# Novel insights into the structure and evolution of the human SAGA complex by affinity-ligand purification

**DOI:** 10.1101/2025.07.31.667873

**Authors:** Mylène Damilot, Thomas Schoeps, Laszlo Tora, Patrick Schultz, Luc Lebeau, Gabor Papai, Adam Ben-Shem

## Abstract

Human SAGA is a 20-subunit complex that stimulates transcription and is essential for development. The most prominent addition to SAGA in metazoans compared to yeast is a 150kDa splicing-factor module (SPL). SPL is also a part of the U2snRNP but its role in SAGA is elusive, partially due to absence of high-resolution structural information regarding its incorporation into the complex. In yeast, subunit TAF5 and TAF6 of SAGA are shared with the general transcription factor TFIID. In metazoans, gene duplication created proteins that occur only in SAGA (TAF5L and TAF6L) or in TFIID (TAF5 and TAF6). What functions of SAGA benefit from this protein specialization is unclear. Here we report the structure of endogenous human SAGA purified via an affinity-ligand from cells that were not disturbed by any genomic engineering tools such as CRISPR-Cas9. Our work reveals the high-resolution structure of SPL and of the TAF6L HEAT repeat domain that provides the SPL with a docking surface. We elucidate how SPL and the HEAT repeats are incorporated into SAGA. We find multiple major differences between TAF6L/TAF5L and the canonical paralogues that are directly implicated in structural re-arrangements required to accommodate SPL. Furthermore, SPL binding to SAGA is very different and occupying much less interaction surface than to U2snRNP. However, the two cases still share similar sequences in a helix that is deeply inserted into the SPL. The seemingly weaker interaction of SPL with SAGA raises the possibility that SAGA serves to relay this module to the splicing machinery. Our structure also suggests mutations that could uncouple SPL from SAGA to further interrogate the role of this module.

## Introduction

Transcription regulation controls many cell decisions including growth, differentiation and response to external cues. DNA-sequence-specific transcription factors recruit co-activators/repressors, many of them are large multi-protein complexes, to stimulate or inhibit transcription by altering chromatin structure. SAGA is such a large complex that modulates chromatin structure via two enzymatic activities: histone acetylation (HAT) and deubiquitination (DUB)[1].

Human SAGA regulates both transcription initiation and elongation and is essential for normal embryonic development [2]. It also has many non-histone targets and thus impacts gene expression at multiple levels from transcription to protein stability [3, 4]. The ∼1.6 MDa complex comprises 20 subunits organized functionally and structurally in five modules: a histone acetyltransferase (HAT) module, a histone deubiquitinase (DUB) module, the TRRAP subunit that serves as a docking surface for transcription activators that recruit SAGA to enhancers, a metazoan-specific 150KDa splicing factors module (SPL) with an enigmatic role and a central module that scaffolds the complex by physically connecting to all other modules. Cryo-electron microscopy (EM) maps of human and yeast SAGA describe well the core and TRRAP parts but do not reveal at high-resolution the HAT, DUB or Splicing modules [5, 6].

GCN5/PCAF is the catalytic subunit of the SAGA HAT module. This subunit is also incorporated into the essential 10-subunit co-activator ATAC, where the HAT module is identical to that of SAGA with the exception of one subunit being replaced by a homologue. SAGA and ATAC are regulating distinct sets of genes [7].

In yeast, the SAGA core module shares with the TBP depositing complex TFIID five subunits: Taf5, 6, 9, 10, and 12. In metazoans, gene duplication created TAF6 and TAF5 paralogues that are specific to SAGA, namely TAF5L and TAF6L. It is currently poorly understood why gene duplication was necessary. What specialized roles do the differences between TAF5L/TAF6L and their yeast or TFIID counterparts play? Are these roles related to the most prominent metazoan-specific feature of SAGA, the splicing module? Answering these questions is hindered by the low-resolution description of the splicing module as well as its docking site on SAGA.

In line with their pivotal role in transcription regulation many co-activators are implicated in human disease, notably cancers. For example, SAGA can be recruited to chromatin by the c-MYC oncoprotein whose deregulation leads to unfavorable variations in the expression of target genes in the vast majority of cancers. The HAT activity of SAGA is necessary for the maintenance of MYC oncogenic transcription program in many tumors [8]. Given the involvement of SAGA in human pathology there is an urgent need to characterize the composition, interactome, activity and structure of endogenous SAGA in cell lines derived from tumors or other human diseases. However, this remains a largely untapped source of information, due to the difficulty in purifying these complexes. Although affinity-tags fused to endogenous subunits of SAGA via CRISPR/cas9 engineering could alleviate this issue, many cell-lines are not amenable to this technology. Moreover, the efforts that such tagging entails impede large comparative studies where SAGA from several different cell-lines are required. Furthermore, optimized purification schemes are required due to the very low abundance of these complexes in cells. For example, a recent study reported only a few picomoles of isolated SAGA from 30 liters of HeLa cell culture [5].

Affinity-ligands, small molecules that bind SAGA and can be coupled to a solid support (e.g. agarose or magnetic beads) offer an attractive alternative to affinity-tags because they can be applied to many different cell-lines. Several small molecules/inhibitors are available that target with high affinity and selectivity the active site of SAGA or its “reader” domains that stabilize interactions with chromatin through recognizing specific histone modifications. However, so far, such molecules were not exploited for developing a purification scheme that yields pure and active SAGA, or any other low-abundance co-activator, let alone enables their structure determination.

In this study, we obtained pure and highly active endogenous human SAGA from two different unmodified cell lines, K562 and HeLa, by employing an affinity-ligand. This compound is composed of an inhibitor for the bromodomain of the enzymatic subunit GCN5/PCAF in the SAGA HAT module and is coupled to desthiobiotin. We introduced several innovations in the design of the affinity ligand as well as in the purification scheme that was adapted to low abundance complexes. Using this new method, we determined the structure of the purified SAGA by cryo-EM and revealed for the first time at high-resolution the splicing factor module in SAGA and as well as the TAF6L HEAT repeats which form the docking site for the splicing and HAT modules. We elucidated in detail how the splicing module and TAF6L HEAT repeats anchor into the co-activator complex. We find that the major deviations in TAF6L with respect to the canonical paralogue are required for incorporating the splicing module. Comparing the docking of this module in SAGA and in the spliceosome, suggests that SAGA serves to relay the splicing module to its assembly in the spliceosome. Our results could potentially guide future endeavors to characterize multi-protein complexes from many medically important sources using affinity-ligands.

## Results

### Design of the affinity ligand and the purification scheme

Low abundance co-activators, such as SAGA are present at a concentration of roughly 1 nM in nuclear extracts. On the other hand, the majority of inhibitors for catalytic enzymes or for histone modification “readers” harbored by co-activators have unfortunately a K_d_ higher than the very low nanomolar range. Hence, the first obstacle that we needed to overcome, in order to make the use of most available inhibitors feasible, was to develop a method for concentrating nuclear extracts. Our experience in yeast and human cells showed that differential precipitation using a high-molecular-weight polyethylene glycol (PEG) is a simple technique to achieve that goal (Fig. 1). Indeed, due to PEG tendency to precipitate first the larger molecules, this technique can also be considered as a real first purification step because it increases the proportion of large complexes in the sample with minimal losses [9]. We worked with nuclear extract derived from 3 L of K562 or 4.5 L of HeLa S3 cells in suspension due to sample requirements for high-resolution structural studies. However much smaller volumes can be used for other applications since PEG precipitation procedure can be easily adapted to practically any volume of nuclear extract.

**Figure 1:**
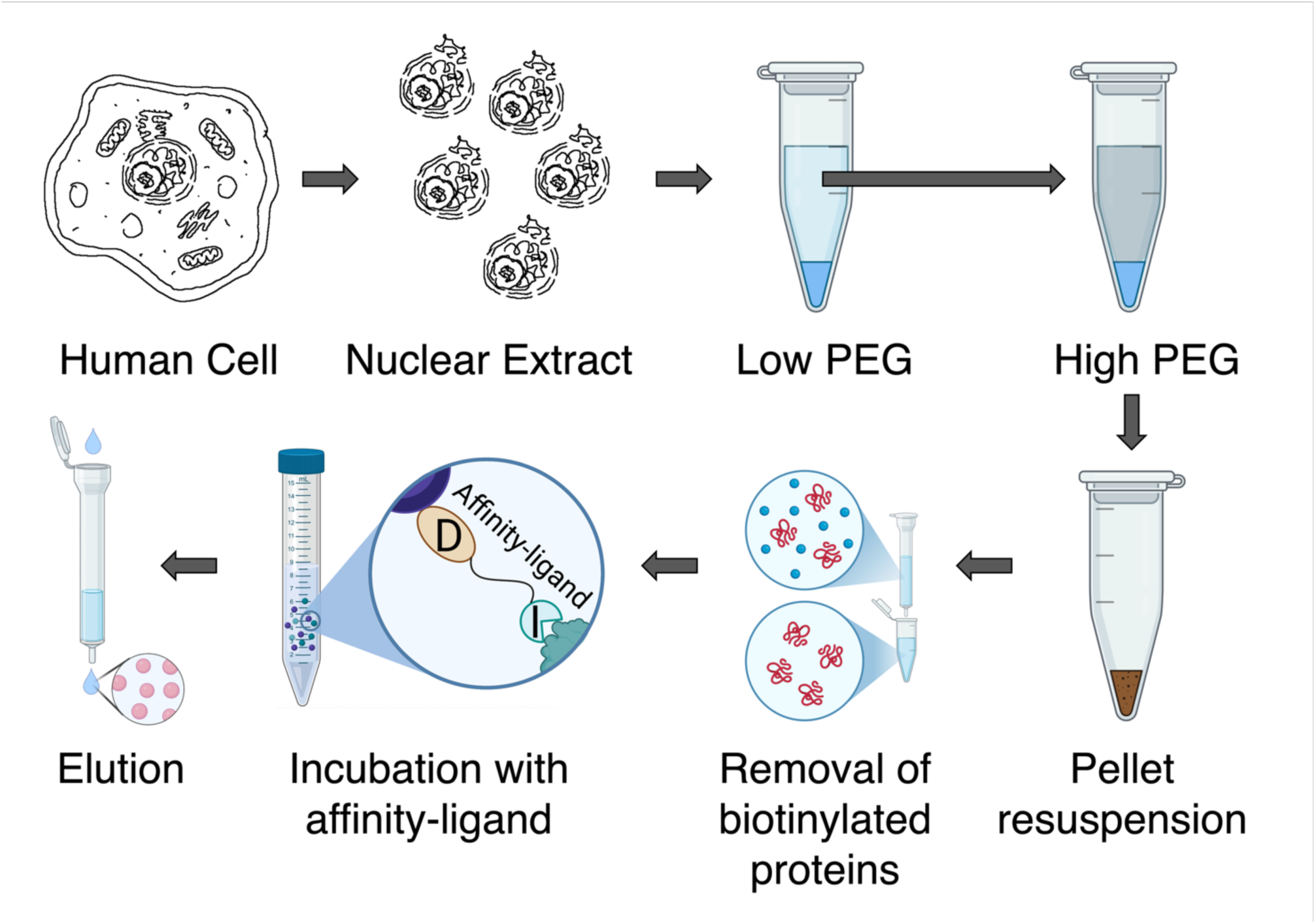
Purification scheme of SAGA and ATAC using affinity-ligand. Schematic overview illustrating the stepwise purification strategy adapted to low-abundance complexes employed to isolate SAGA and ATAC (Low and High PEG: precipitation with low and high concentration of polyethylene glycol; I: inhibitor; D: desthiobiotin). Figure was created with help of BioRender (https://BioRender.com).

We then screened the literature for high affinity and selectivity binders for SAGA. We chose eventually GSK4027 [10] which targets the bromodomain, a “reader” that recognizes acetylated lysines, in the catalytic HAT subunit GCN5/PCAF with a K_d_ = 1.4 nM (Supplemental Fig. 1a). This compound is highly selective to GCN5/PCAF, presenting much lower affinity to other bromodomains. One drawback of using GSK4027 as a ligand is that it cannot distinguish between the two human complexes that harbor GCN5/PCAF namely ATAC and SAGA. However, we posited that due to the predicted size difference between SAGA (1.68 MDa) and ATAC (0.9 MDa), *in-silico* classification of the cryo-EM dataset would be able to separate the two complexes.

To mediate the binding of GSK4027 to a solid matrix we chose to conjugate it to desthiobiotin which can be attached to streptavidin beads (K_d_ = 10^-11^ M) and efficiently eluted under native conditions with biotin (K_d_ = 10^-16^ M). From X-ray crystallographic data on GSK4027 bound to the human GCN5/PCAF bromodomain, the 4-position on the pendant phenyl ring seems to point outside the binding pocket [10]. This is indeed confirmed by results obtained by Bassi *et al*. who developed a GSK4027-based PROTAC approach for modulating GCN5/PCAF immune cell function [11]. This position was thus selected for derivatization with desthiobiotin while preserving binding to the bromodomain. Consequently, we designed conjugates **1a** and **1b** as ligands for immobilization of SAGA/ATAC on streptavidin beads and purification by affinity chromatography (Supplemental Fig. 1b). The GSK4027 and desthiobiotin moieties are connected through an oligoethylene spacer of variable length to modulate access of each ligand to its respective target: GSK4027 to the bromodomain and desthiobiotin to streptavidin beads and alleviate possible steric hindrance.

### Synthesis of the affinity ligands 1a and 1b

Prior to our work the synthesis of GSK4027 was achieved by Humphreys *et al*. in 7 steps [10]. The compound was obtained as a mixture of four enantiomers and final separation of the racemate by preparative chiral HPLC was necessary to get the desired single enantiomer in an overall yield of 1.2%. In this study, we developed a stereo-controlled synthetic route to prevent the formation of mixed isomers. This approach aligns with a methodology previously established by Steijvoort *et al [12]* involving a Pd-catalyzed stereoselective C5(sp3)-H arylation of 1-Boc-3-(picolinoylamino)piperidine with an aryl iodide [12]. The synthesis of the GSK4027 scaffold (compound 7) is a six-step process with an overall yield of 7.6% (Supplemental Fig. 1c). Two additional steps are required to for the target compounds 1a and 1b. The full experimental details are provided in the Material section.

### Purification of active SAGA/ATAC using affinity-ligand

Nuclear extracts from cell lines K562 or HeLa S3 are first treated with very low (∼1%) concentration of PEG 20,000, to deplete the extracts from large particles such as membrane parts and vesicles. A higher concentration (∼5%) of PEG is then used to pellet large multi-protein complexes including SAGA and ATAC. The pellet is suspended in a small volume that is found to contain almost all SAGA/ATAC particles but only 30-40% of the total starting protein content (Fig. 2a-c). Hence the PEG precipitation step concentrates the sample by roughly ∼20 times with minimal loss and at the same time increases the proportion of the target complexes. We obtained similar results with other large complexes [9].

**Figure 2:**
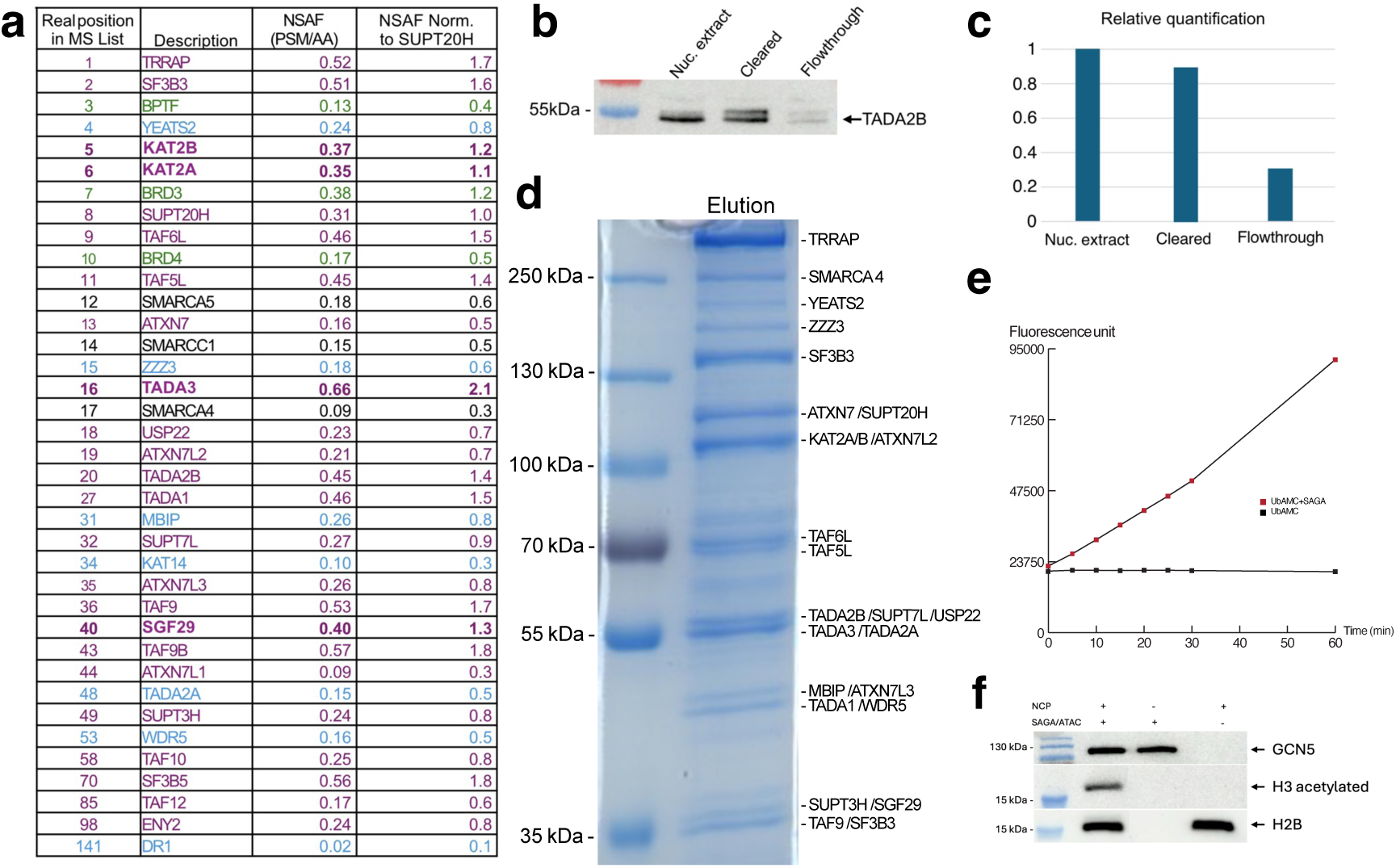
Purification of SAGA/ATAC using affinity-ligand with long linker. **a**, Proteomic analysis of the purified SAGA and ATAC complexes. For each identified protein the table shows the NSAF value calculated from the Peptide Spectrum Matches (PSM) divided by the number of amino acids, and the normalization of the NSAF value to SUPT20H as rough estimation of stoichiometry. SAGA subunits are colored purple, ATAC subunits are colored blue, purple bold represents the common subunits between SAGA and ATAC, and co-purified bromodomain containing proteins are in green. **b**, Western blot analysis of different purification steps detecting the TADA2B subunit of SAGA. **c**, Relative quantification of the detected bands in panel b. d, Colloidal Coomassie blue stained SDS-PAGE of the purified SAGA and ATAC complexes, the identity of indicated bands was verified by mass spectrometry. **e**, Deubiquitination activity of the purified SAGA on the fluorogenic Ubiquitin-amino methyl coumarin (Ub-AMC) over time. **f**, Acetyltransferase activity of the purified SAGA and ATAC complex on Nucleosome Core Particles (NCPs). Acetylation of histone was detected by a pan-acetyl-lysine antibody. Anti-GCN5 and H2B antibodies were used as loading control of SAGA/ATAC and NCPs respectively.

We analyzed the eluted sample derived from wild type K562 cells by SDS-PAGE and mass spectrometry (MS) (Fig. 2a and d). The isolated complex is ∼90% pure as estimated by a polyacrylamide gel that also shows all known subunits of SAGA and ATAC with the exception of the very small ones (Fig. 2d). Similarly, MS analysis identifies all predicted subunits of SAGA and ATAC (Fig. 2a). Moreover, roughly 60% of the total MS signal - estimated by summing all peptide spectrum matches (PSMs) – corresponds to subunits of SAGA and ATAC complexes. This high value is close to that obtained by tandem-tag affinity purification of low-abundance complexes yielding high resolution cryo-EM structures [9]. We note that purity estimation by MS is always much lower than SDS-PAGE due to its high sensitivity. Among the 20 subunits with highest signal in the MS data, only six are not attributed to SAGA/ATAC (Fig. 2a). Three of the six are bromodomain-containing proteins (BPTF, BRD3, BRD4) and probably present some specific affinity for the GSK4027 ligand. However, compared to SAGA, these are small proteins that will be easily removed in the cryo-EM data analysis and what is more, their stoichiometry in the MS table is lower than SAGA subunits.

SDS-PAGE and MS analysis of eluted sample derived from HeLa S3 cells showed similar results demonstrating the versatility of our purification method that can be applied to different unmodified cell lines (Supplemental Fig. 2). We elected to continue our investigation only with the K562 cells because the SAGA concentrations seemed higher in this source.

To understand the importance of the linker that connects desthiobiotin to GSK4027 we compared purification that employs ligand 1a (i.e., with a linker made of three ethylene glycol units) instead of 1b. We find that linker length has a profound and critical effect on the quality of the purification. Employing the conjugate with the shorter linker captures only a small fraction of SAGA/ATAC on the beads and yields a highly contaminated sample (Supplemental Fig. 3).

To further demonstrate the quality of SAGA/ATAC purified by employing our novel affinity-ligand we performed activity assays. First, we tested the capacity of SAGA to cleave immediately after a ubiquitin using the fluorogenic substrate, ubiquitin-amino methyl coumarin (Ub-AMC) [13]. As Figure 2e demonstrates, SAGA is highly active in cleavage of Ub-AMC, releasing the AMC moiety from ubiquitin and thus giving rise to a strong fluorescence signal. Second, we incubated SAGA/ATAC with nucleosomes in the presence of acetyl-CoA and followed acetylation of histones using an antibody that recognizes acetylated-lysines. It seems that SAGA/ATAC are highly active in acetylating histone H3 (Fig. 2f). Clearly, we cannot exclude the possibility that only one of the two complexes is active, but this seems unlikely as both were purified together, harbor a nearly identical HAT module, and appear intact in MS and SDS-PAGE.

Using PEG clearance and concentration of nuclear extracts as a starting point to a purification that employs an affinity-ligand conjugated via a long (six-ethylene glycol units) linker to desthiobiotin, we are able to isolate highly pure and active native tag-less co-activators in relatively large amounts.

### Structure of SAGA – overview

We elucidated the structure of SAGA at 2.4-3 Å resolution (Supplemental Fig. 4). Our structure of human SAGA supports earlier descriptions of the complex. Human SAGA has a three-lobed architecture. One lobe consisting mainly of the TRRAP activator-platform subunit, a second includes a Y-shaped splicing module and the third, core lobe, is orchestrated by the TAF5 WD40 domain and contains a deformed octamer of histone folds (Fig. 3). As noted by others the major deviation between human and yeast SAGA, apart from the additional splicing module, is the rotation of the TRRAP subunit around the core by nearly 80° [5]. Furthermore, the interaction between the core and TRRAP is much more extensive in the human case where most core subunits contribute to the contact compared to only three in yeast [6].

**Figure 3:**
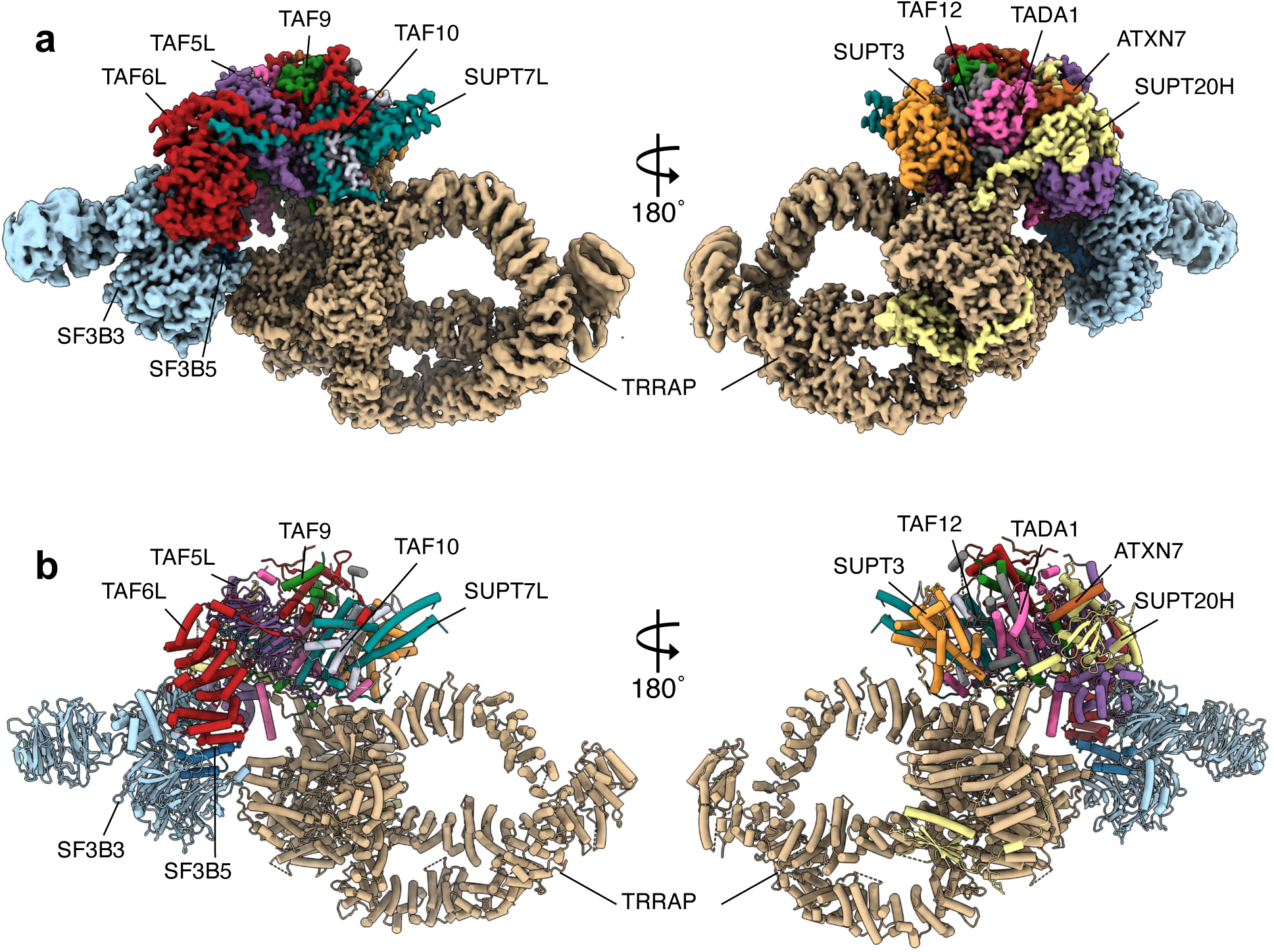
Structure of the human SAGA complex purified by long-linker affinity-ligand. **a**, Two views of the composite cryo-EM map of human SAGA **b**, Corresponding views of the atomic model of the SAGA complex.

Two additional lobes appear in the yeast 3D maps of SAGA at very low resolutions and correspond to the HAT and DUB modules that seem to maintain some contact between themselves as is suggested by cross-linking data as well [14]. In the 3D maps of human SAGA, we find no trace of HAT and DUB although mass-spectrometry strongly indicates that they are attached to SAGA and moreover the complex is purified via a ligand that recognizes the HAT module. This would suggest that the flexibility of these modules in the human case is enhanced compared to yeast. Indeed, in yeast the DUB module associates with the core through the N-terminal domain (NTD) of TAF5 and the HAT seems to have some ties to Spt7 [15]. In the human counterpart however TAF5L NTD is translocated to a different position to vacate space for the splicing module [5]. Similarly, human Spt7 (SUPT7L) is shorter by 600 residues than its yeast homologue and has much weaker, if any, contact with the HAT module. Hence the loss of contacts with the core could underlie an enhanced autonomy or flexibility for the HAT and DUB modules.

### The splicing module in SAGA

Earlier studies of SAGA could not resolve the structure of the 150kDa splicing module, composed of the factors SF3B3 and SF3B5, that is absent in yeast SAGA. They also failed to define in high resolution the TAF6L HEAT repeats, which are of crucial importance in SAGA because they are the docking site of the HAT and splicing modules and provide their connection to the core. Therefore, a high-resolution understanding of how the splicing module docks on SAGA and how the TAF6L HEAT repeats are incorporated into the core are lacking. The structure of endogenous human SAGA purified by affinity-ligand can fill this gap in our understanding.

We obtained maps with a resolution of roughly 3 Å on the splicing factors and TAF6L HEAT repeats which allowed us to analyze their interaction at the level of sidechains (Fig. 4a, b and c). The SF3B3 and SF3B5 factors also associates with protein SF3B1 in the SF3B complex, an intrinsic component of the functional U2snRNP, part of the mRNA splicing machinery. We could therefore compare the TAF6L-SPL contacts in SAGA to the association of SPL with SF3B1 in the SF3B complex (Fig. 4d and e). We find that the partners of the splicing module, namely SF3B1 and TAF6L HEAT repeats, form a right-handed and left-handed super helix, respectively (Fig. 4d and e). This inherent difference in their architecture results in little overlap between the positions of their HEAT repeat helices with respect to SPL. The networks of interactions in the two cases are different in general and have little in common. However, two similarities stand out. First, both TAF6L HEAT repeats and SF3B1 interact with the same face of SPL and are therefore mutually exclusive. Second, the last HEAT repeat helix of both SF3B1 and TAF6L are both located in the same place with respect to SPL and contribute considerably to the interaction with SF3B3 and SF3B5 (Fig. 4f). Furthermore, we find that the tip of this helix, namely the last seven residues, have very similar sequences in both cases (DSLATRF in TAF6L, DALIAHY in SF3B1) and establish similar interactions with SPL (Fig. 4g, h).

**Figure 4:**
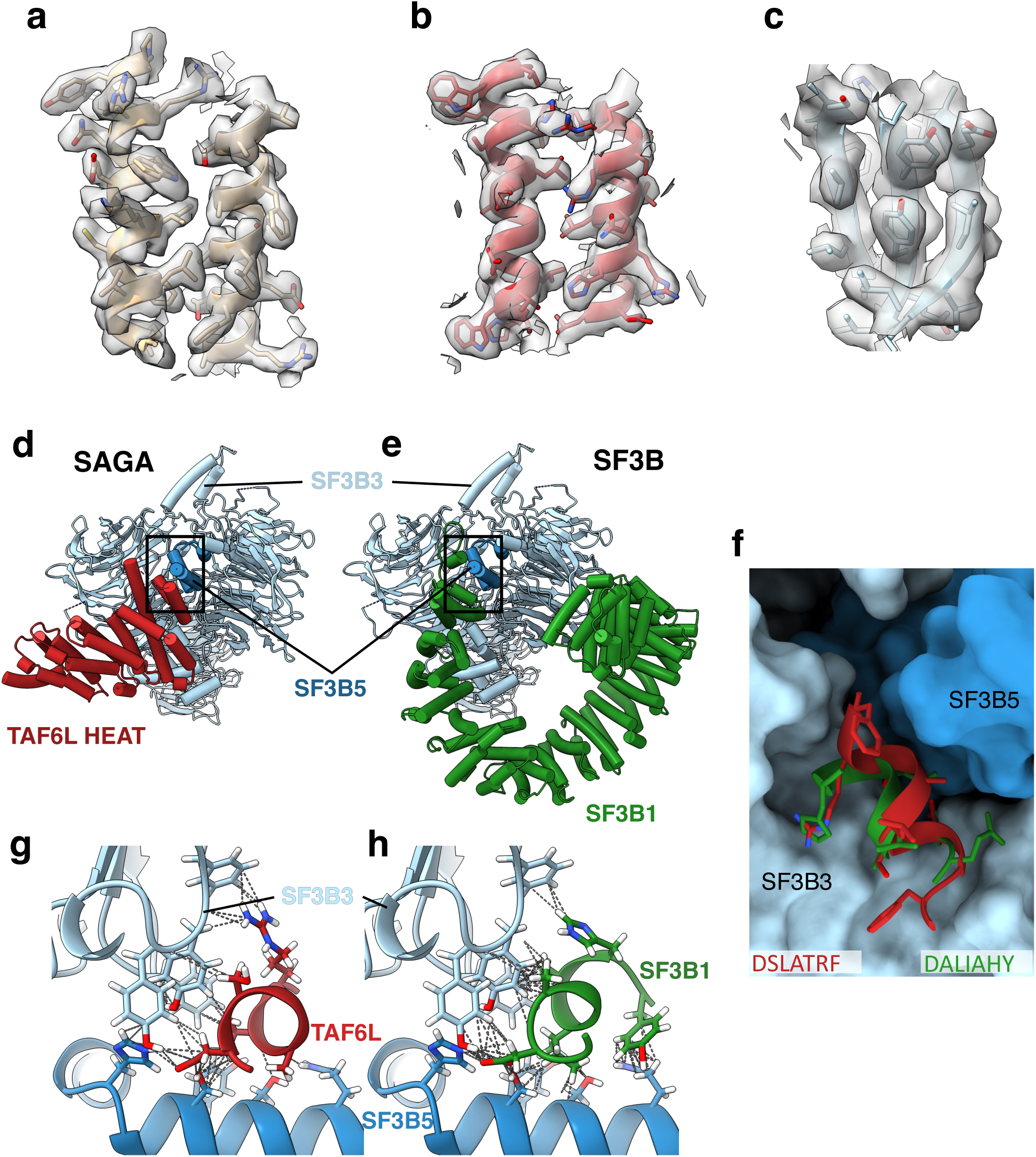
Splicing factor SF3B interaction with SAGA complex. **a, b**, and **c**, Representative cryo-EM densities showing the map quality in the TRRAP, TAF6L HEAT repeats and SF3B regions, respectively. **d**, Interaction between SF3B3 (light blue)-SF3B5 (dark blue) splicing module with TAF6L HEAT repeats (red) in human SAGA. **e**, Interaction between SF3B3 (light blue)-SF3B5 (dark blue) splicing module with SF3B1 (green) in SF3B. **f**, Close-up view of the homologous last helix in TAF6L HEAT repeats (red) and SF3B1 (green) interacting with a surface formed by SF3B3 (light blue surface) and SF3B5 (dark blue surface). **g**, Interaction between the last helix in TAF6L HEAT repeats (red) and SPL (SF3B3 – light blue and SF3B5 – dark blue). The contacts are shown with dotted lines. h, Interaction between the last helix in SF3B1 HEAT repeats (green) and SPL (SF3B3 – light blue and SF3B5 – dark blue). The contacts are shown with dotted lines.

It is noteworthy that in SF3B1 this last helix is continued by a 30 amino acid linker that is inserted deep into a cleft between SF3B5 and SF3B3, interacting with both proteins, and probably playing an important role in stabilizing the interaction in SF3B. Overall we find that the buried surface involved in the interaction between TAF6L and SPL in SAGA is only half of that between SF3B1 and SPL in SF3B (1500 Å^2^ vs 3000 Å^2^). This suggests that SAGA binds SPL much weaker than the functional U2snRNP.

Another noteworthy observation regarding the SPL is the location of the seven-bladed β-propeller (BP) domain known as BPB (residues 472-770). We find that the position of this domain with respect to the rest of the module is shifted by 20 Å when compared to the crystal structure of the SF3B complex [16]. This position is similar to the position of BPB within the spliceosome [17]. This suggests that a similar conformation change accompanies loading of SPL into SAGA or spliceosome.

### Incorporating the TAF6 HEAT repeats - importance of gene duplication creating SAGA specialized genes

TAF6L HEAT repeats are a structural hub that coordinates the HAT and splicing module and connects them to the core. In yeast the TAF5 NTD and the SPT7 NTD play an important role in stabilizing the TAF6 HEAT repeats [6, 18]. However, in the human case TAF5L NTD is displaced by SPL and SUPT7L NTD is just a short linker that only weakly, if at all, contributes to binding the HEAT repeats. These changes were probably required for adding the splicing module and affording greater autonomy to the HAT. But what evolutionary adjustments made these changes possible while still maintaining TAF6L HEAT repeats well docked into the core?

The answer lies in gene duplication events, creating specialized genes for SAGA rather than genes shared with other complexes as is the case in yeast. The most significant difference in sequence alignment between TAF6 and TAF6L lies in the linker that connects the histone fold with the HEAT repeat (Supplemental Fig. 5a). En route, this linker complements the first of the TAF5/TAF5L WD “propeller” blades with two β-strands. In yeast and in human TAF6, this linker then continues with ∼60 additional amino acids before forming the first HEAT repeat. In the SAGA-specialized TAF6L only 6 residues separate the end of the β-strands and the base of the first HEAT repeats helix (Supplemental Fig. 5b). Hence the interaction of the linker with the TAF5L propeller forms an anchor point that strongly constrains the position of the TAF6L HEAT repeats. The second important adjustment occurs in the TAF5L helix that leads into the first WD propeller blade. In human TAF5L, this helix is shifted by 12 Å and tilted by 92° relative to yeast Taf5, and by 16 Å and 106° relative to human TAF5 (Fig. 5a). This is due to the fact that this helix is connected to the first propeller blade via a much shorter linker in the TAF5L (5 residues compared to 12 in both yeast and human TAF5). As a result, this helix in human SAGA runs anti-parallel to the first HEAT repeat, with which it forms a three-helix bundle that has the most important contribution to the positioning and stability of the TAF6L HEAT repeats domains (Fig. 5b). Strikingly, in human TFIID, at the same position with respect to the TAF6 HEAT repeats in lobe C, we find a helix attributed to subunit TAF8 that runs parallel to the repeats and forms a three-helix bundle with the first repeat, very similar to the TAF5L helix (Fig. 5c). Thus, a common solution to the requirement of positioning the TAF6L/TAF6 HEAT repeats is applied both by SAGA and TFIID albeit with different subunits, namely TAF5L and TAF8 respectively.

**Figure 5:**
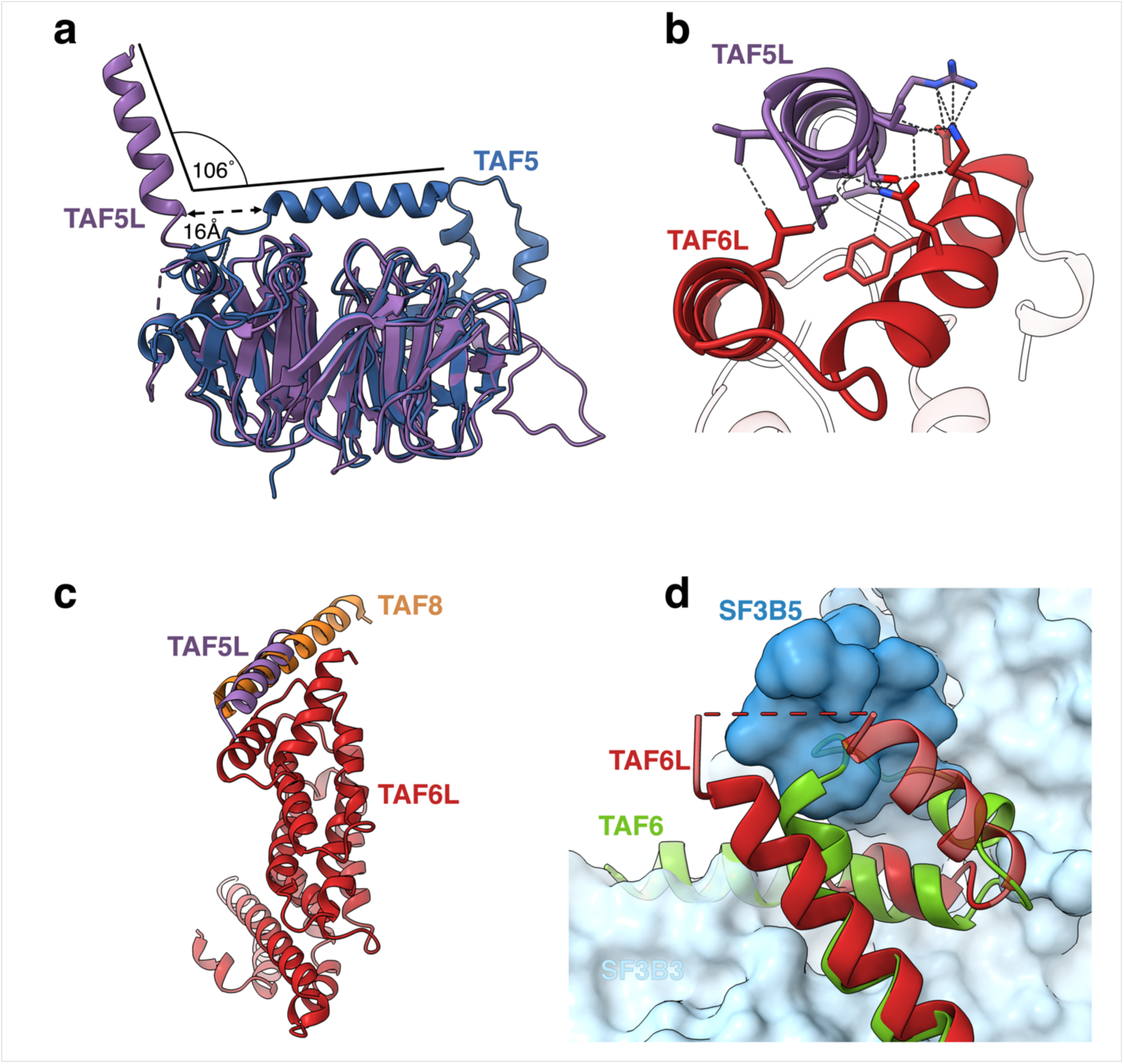
SAGA-specific adaptations enable SPL incorporation. **a**, Comparison of **human** TAF5 (in TFIID – blue) and TAF5L (in SAGA – violet). The TAF5L helix interacting with TAF6L HEAT repeat domain in SAGA is displaced compared to its counterpart in TAF5. **b**, The same TAF5L helix forms a 3-helix bundle with the HEAT repeat domain of TAF6L (red) in SAGA. The contacts are shown with dotted lines. **c**, TAF8 (orange) in TFIID interacts similarly in TFIID with TAF6 as TAF5L (violet) with TAF6L (red). TAF6 HEAT repeats were aligned on TAF6L HEAT. Only TAF6L repeats are shown. **d**, The last HEAT repeat of TAF6L (red) deviates from the one of TAF6 (green) to allow the incorporation of the splicing module (SF3B3 – light blue surface and SF3B5 – dark blue surface) into SAGA. The last HEAT repeat of TAF6 clashes with both SF3B3 and SF3B5.

In addition to the long linker connecting the histone fold and HEAT repeats, the main structural difference between TAF6 and TAF6L appears in the last HEAT repeat which assumes a very different angle in TAF6 and ends with a much longer helix. Superposing the TAF6 and TAF6L HEAT repeats shows that the TAF6 repeats would clash with SF3B3 (Fig. 5d) and of course the multitude of interactions formed by the tip of the last helix in TAF6L and SPL (Fig. 4f) would not be reproduced by the completely different sequence in TAF6.

Hence gene duplications that created proteins specialized for a role in SAGA were crucial for the structural re-arrangement that is required for adding the splicing module and, probably, augmenting HAT autonomy.

## Discussion

We present in this paper the first example, as far as we are aware, for an isolation based on small compound affinity-ligand yielding an intact, active and highly pure low-abundance multi-protein complex suitable for high resolution structure determination. We could employ our SAGA isolation method with two different cells lines, namely K562 and HeLa S3, demonstrating the applicability of this concept for different sources. We further show the utility of this method and the quality of the sample by generating the highest resolution structure ever obtained of the human SAGA complex. In addition to the choice of the affinity-ligand, we ascribe this success to two innovations we introduced, namely a long linker to conjugate the affinity ligand to desthiobiotin and the use of PEG precipitation to clear the nuclear extract and concentrate it.

The structure of full GCN5/PCAF, let alone the holo-HAT module of SAGA/ATAC, has not been elucidated so far and therefore it is unknown where the bromodomain target of GSK4027 is situated within this module. We can only speculate that the affinity ligand with the long linker allows to reach the bromodomain without SAGA/ATAC clashing with the solid matrix. The bromodomain of GCN5/PCAF binds acetylated lysines on the N-terminal tail of histone H3, which is a long, flexible, thread of roughly 40 amino acids. It is therefore possible that the bromodomain is not located on the surface of the HAT module but rather situated in a cavity where it is still accessible to the flexible H3 tail. Such a cavity would indeed necessitate a long linker for GSK4027 to infiltrate inside without bringing SAGA/ATAC too close to the matrix.

The possibility to efficiently isolate intact and highly pure SAGA from practically any cell type leads us to suggest that purification of other co-activator complex is feasible provided that a suitable affinity-ligand is available. Isolation of these complexes from different organisms or stages of development, from various organs or from different pathologies could be crucial for elucidating the interactome of co-activators and their place in the pathways that lead to disease and differentiation. For the specific case of SAGA which we addressed here, we note for example that spinocerebellar ataxia 7 is an adult-onset neurodegenerative disorder caused by the expansion of a CAG repeat sequence located within the coding region for SAGA DUB subunit ATXN7. The underlying molecular mechanism for the disease is not fully elucidated [19]. Isolating SAGA from cells of patients (or animal models) at different ages and measuring its deubiquitination activity could shed new light on the physiopathology of this disease. Another instance is presented by regulatory T-cells that attenuate the immune response, where FOXP3 is a master regulator of multiple pathways crucial for their function. SAGA plays a critical role in controlling FOXP3 and therefore understanding the SAGA-FOXP3 interaction in these cells could help to better manage autoimmunity or cancer [20].

Our work reveals for the first time the high-resolution structures of the 150KDa splicing factor module on SAGA and the TAF6L HEAT repeats, as well as the molecular details of their incorporation into SAGA. Possibly the very gentle nature of our purification scheme and the fact that we could employ wild type cells whose genomes were not manipulated by CRISPR/Cas9, allowed us to determine maps with overall higher resolution and in particular better description of this peripheral region of SAGA. We could therefore compare in detail the interaction of SPL with TAF6L HEAT repeats in SAGA to their docking on SF3B1 in SF3B. Although both TAF6L and SF3B1 associate with the same face of SPL, the interaction networks are very different, apart from the last helix in both proteins that is positioned in the same place with respect to SPL. The last helices have very similar sequences at their last two turns and contributes significantly to the contact with SF3B3 and SF3B5.

At least in the case of TAF6, and possibly also of TAF5, the most important structural differences between the SAGA-specialized protein (TAF6L) and the canonical versions in TFIID or in yeast, are directly related to the structural re-arrangements required to accommodate the splicing module. These differences include a linker, shorter in TAF6L by ∼50 aa compared to TAF6, that constrains the position of the TAF6 HEAT repeats, a shorter linker in TAF5L that positions a helix to form a helix bundle with the first HEAT repeat, and a completely different last two-helices of the TAF6L HEAT repeats where the TAF6 version would clash into the SPL and would fail to present important residues for binding the SPL. This suggests that to a considerable extent the gene duplication events in metazoans occurred to enable the incorporation of the splicing module which is the most prominent addition to metazoan SAGA compared to yeast. This fact suggests that the role of the splicing module in-vivo is of considerable importance, although elusive. Our work proposes where mutations can be introduced to uncouple SPL from the HEAT repeats and SAGA so that its role could be studied. Eliminating the last two HEAT repeats of TAF6L or possibly even just the last helix should considerably undermine the SAGA contact with SPL.

Importantly, we find that the buried surface in the SAGA interaction with SPL is only half that in the SF3B counterpart. The known correlation between buried surface and binding affinity [21] clearly suggests that SPL is associated much stronger with SF3B than with SAGA. Significantly contributing to this firmer interaction is the 30 amino acids linker in SF3B1, without any parallel in SAGA, that is inserted deeply into the cleft between SF3B3 and SF3B5 and forms a very extensive network of contacts with both subunits. We propose that SAGA serves as a relay of splicing factors that transfers SPL for assembly into the pre-spliceosome when SF3B1 (or the entire U2snRNP) are in the proximity of chromatin-bound SAGA during transcription-coupled splicing. Findings in *S. cerevisiae* seem to support this idea. The yeast SAGA physically binds the U2snRNP ATPase PRP5 and modulates its activity in pre-mRNA branch-point proofreading [22]. Interestingly, this activity requires an interplay between SF3B1 and PRP5 [23]. We further suggest that the role of SAGA in relaying SPL to SF3B/U2snRNP could facilitate synchronization between transcription initiation and splicing. It could also limit accumulation of functional U2snRNP in the vicinity of mRNA thus perhaps preventing possible splicing errors.

## Methods

### Synthesis of the affinity ligands

Unless otherwise stated, all chemical reagents were purchased from abcr GmbH (Karlsruhe, Germany) and used without purification. When required, solvents were dried just before use. Thin layer chromatography (TLC) was performed on precoated plates (0.25 mm Silica Gel 60, F_254_, Merck, Darmstadt, Germany). Products were purified by flash chromatography over silica gel (Silica Gel 60, 40-63 μm, Merck, Darmstadt, Germany). NMR spectra were recorded on a Bruker 400 MHz Avance III or Bruker 500 MHz Avance III instrument. ^1^H- and ^13^C-NMR chemical shifts δ are reported in ppm relative to their standard reference (^1^H: CHCl_3_ at 7.26 ppm, CD_2_HOD at 3.31 ppm; ^13^C: CDCl_3_ at 77.0 ppm, CD_3_OD at 49.0 ppm). Splitting patterns are assigned s = singlet, d = doublet, t = triplet, q = quartet, quin = quintet; br. = broad signal. IR spectra were recorded on a FT-IR Nicolet 380 spectrometer in the ATR mode and absorption values v are in wave numbers (cm^-1^). Mass Spectra (MS) were recorded on an Agilent Technologies 6520 Accurate Mass QToF instrument, using electrospray ionization (ESI) mode. Mass data are reported in mass units (*m/z*).

#### *tert*-Butyl (3R)-3-(pyridine-2-carbonylamino)piperidine-1-carboxylate (2)

To a solution of (R)-3-amino-1-*N*-Boc-piperidine (250 mg, 1.25 mmol) and 2-picolinic acid (189 mg, 1.53 mmol) in anhydrous CH_2_Cl_2_ (20 mL) under inert atmosphere was added EDC.HCl (359 mg, 1.87 mmol) and 4-DMAP (15 mg, 0.12 mmol). The reaction mixture was stirred at rt until the starting piperidine was no more detected by TLC (*ca*. 2.5 h). Then it was washed with saturated aqueous NaHCO_3_, and brine. The organic layer was dried over MgSO_4_, filtered and reduced under vacuum. The crude residue was purified by flash chromatography on silica gel (PE/Et_2_O: 20/80) to yield compound **1** (374 mmol, 98%) as a white powder. Analyses were consistent with the literature data.

#### *tert*-Butyl(3*R*,5*R*)-3-(4-(methoxycarbonyl)phenyl)-5-(picolinamido)piperidine-1-carboxylate (3)

Compound **3** was obtained from **2** according to the protocol described by van Steijvoort *et al. [12]*.

#### Methyl 4-((3*R*,5*R*)-5-(picolinamido)piperidin-3-yl)benzoate (4)

Compound **3** (1.231 g, 2.80 mmol) was stirred in CH_2_Cl_2_/TFA 9:1 (25 mL) at rt for 2 h. Volatile was removed under vacuum, the residue was stirred in CH_2_Cl_2_/H_2_O 1:1 (20 mL) and pH was set at 12 with NaOH 1 N. The organic layer was separated, and the aqueous layer was extracted twice with CH_2_Cl_2_. The combined organic layers were dried over MgSO_4_, filtered and reduced under vacuum to yield compound **4** (0.885 g, 92%) as a slightly yellow powder. ^1^H-NMR (CDCl_3_, 500 MHz) δ 8.49 (dd, *J* = 4.8, 0.8 Hz, 1H), 8.16 (d, *J* = 7.8 Hz, 1H), 7.94 (m, 3H), 7.80 (td, *J* = 7.7, 1.7 Hz, 1H), 7.38 (ddd, *J* = 7.6, 4.7, 1.2 Hz, 1H), 7.26 (d, *J* = 8.3 Hz, 2H), 4.25 – 4.07 (m, 1H), 3.86 (s, 3H), 3.42 (m, 1H), 3.17 (m, 1H), 2.90 (m, 1H), 2.62 (dd, *J* = 12.6, 11.2 Hz, 1H), 2.53 (dd, *J* = 12.2, 10.8 Hz, 1H), 2.35 (m, 1H), 1.68 (q, *J* = 12.1 Hz, 1H). ^13^C-NMR (CDCl_3_, 101 MHz) δ 166.79; 163.64; 149.64; 148.17; 147.87; 137.27; 129.76 (2C); 128.41; 126.94 (2C); 126.10; 122.10; 52.43; 51.9; 51.11; 47.45; 43.83; 38.11. HRMS (ESI) for C_19_H_22_N_3_O_3_ [M+H]^+^, calcd 340.4025, found 340.4032.

#### Methyl 4-((3*R*,5*R*)-1-methyl-5-(picolinamido)piperidin-3-yl)benzoate (5)

Formaldehyde 37% (350 µL) was added to a stirred mixture of compound **4** (990 mg, 2.91 mmol) and acetic acid (35 µL) in MeOH (20 mL) at rt. Then NaBH(OAc)_3_ (1.33 g, 6.27 mmol) was added portion wise over a 20-min period and the mixture was stirred for 1 h before solvent was removed under vacuum. The crude residue was suspended in CHCl_3_ and washed with water at pH 12. The aqueous layer was washed three times with CHCl_3_ and the organic layers were combined together, dried over MgSO_4_, and reduced under vacuum to yield **5** (943 mg, 92%) as a white powder. *R*_f_ 0.5 (CH_2_Cl_2_/MeOH 9:1). ^1^H-NMR (400 MHz, CDCl_3_) δ 8.53 (ddd, *J* = 4.8, 1.7, 0.9 Hz, 1H), 8.19 (dt, *J* = 7.8, 1.1 Hz, 1H), 8.01 – 7.93 (m, 3H), 7.85 (td, *J* = 7.7, 1.7 Hz, 1H), 7.42 (ddd, *J* = 7.6, 4.8, 1.2 Hz, 1H), 7.36 – 7.28 (m, 2H), 4.36 (m, 1H), 3.90 (s, 3H), 3.25 (d, *J* = 13.0 Hz, 1H), 3.12 (t, *J* = 14.6 Hz, 1H), 2.99 (d, *J* = 11.3 Hz, 1H), 2.38 (s, 3H), 2.32 (d, *J* = 15.8 Hz, 1H), 2.07 (t, *J* = 11.2 Hz, 1H), 1.97 (m, 1H), 1.57 (m, 1H). ^13^C-NMR (CDCl_3_, 101 MHz) δ 166.92; 163.68; 149.77; 148.24;147.95; 137.438 129.84 (2C); 128.58; 127.17 (2C); 126.18; 122.22; 61.68; 60.25; 52.00; 46.51; 45.93; 41.49; 37.21. HRMS (ESI) for C_20_H_24_N_3_O_3_ [M+H]^+^, calcd 353.4209, found 353.4212.

#### Ethyl 4-((3*R*,5*R*)-5-amino-1-methylpiperidin-3-yl)benzoate (6)

Picolinamide **5** (100 mg, 0.28 mmol) was stirred in refluxing HCl 12 N for 20 h. The reaction mixture was reduced under vacuum and the residue was coevaporated twice with EtOH. Then it was stirred with acetyl chloride (0.1 mL) in refluxing EtOH (4 mL) for 3 h. Volatile was removed, and the dry residue was suspended in AcOEt and washed with NaOH 2 N. The aqueous layer was extracted twice with AcOEt and the combined organic layers were dried over MgSO_4_ to yield an orange oil that was purified by flash chromatography over silica gel (CHCl_3_/MeOH/H_2_O 9:1:0 to 10:6:1) to afford **6** (49 mg, 66%). *R*_f_ 0.2 (CH_2_Cl_2_/MeOH 9:1). ^1^H-NMR (400 MHz, CD_3_OD) δ 7.98 (d, *J* = 8.4 Hz, 2H), 7.39 (d, *J* = 8.4 Hz, 2H), 4.36 (q, *J* = 7.1 Hz, 2H), 3.13 – 2.90 (m, 4H), 2.38 (s, 3H), 2.18 – 2.08 (m, 1H), 2.05 (t, *J* = 11.9 Hz, 1H), 1.88 (t, *J* = 11.9 Hz, 1H), 1.41 (m, 1H), 1.40 (t, *J* = 7.1 Hz, 3H). ^13^C-NMR (101 MHz, CD_3_OD) δ 167.91, 149.74, 130.82, 130.19, 128.42, 63.13, 62.35, 62.07, 46.12, 42.70, 40.10, 14.60. HRMS (ESI) for C_15_H_23_N_2_O_2_^+^ [M+H]^+^, calcd 263.1754, found 263.1766.

#### Ethyl 4-((3*R*,5*R*)-5-((5-bromo-1-methyl-6-oxo-1,6-dihydropyridazin-4-yl)amino)-1-methylpiperidin-3-yl) benzoate (7)

A solution of DBU (68 µL, 0.46 mmol), 4,5-dibromo-2-methylpyridazin-3(2H)-one (123 mg, 0.42 mmol), and compound **6** (120 mg, 0.42 mmol) in dimethylacetamide (1 mL) was stirred at 120°C overnight in an airtight sealed tube. The reaction mixture was then cooled to room temperature, CHCl_3_ (10 mL) was added and the mixture was washed with NaOH 1 N (4 x 5 mL). The organic layer was dried over MgSO_4_, filtered and reduced under vacuum. The crude residue was purified by flash chromatography over silica gel (AcOEt/EtOH 9:1) to afford **7** (116 mg, 57%). *R*_f_ 0.25 (AcOEt/EtOH 9:1). Analyses were consistent with the literature data [12].

#### 4-((3*R*,5*R*)-5-((5-Bromo-1-methyl-6-oxo-1,6-dihydropyridazin-4-yl)amino)-1-methylpiperidin-3-yl)benzoic acid (8)

Compound **8** was obtained from **7** following the procedure described by Bassi *et al [11]*.

#### *N*-(2-(2-(2-Azidoethoxy)ethoxy)ethyl)-6-((4*R*,5*S*)-5-methyl-2-oxoimidazolidin-4-yl)hexanamide (9a)

Desthiobiotin (100 mg, 0.47 mmol), 2-(2-(2-azidoethoxy)ethoxy)ethan-1-amine^3^ (120 mg, 0.69 mmol), Et_3_N (200 µL, 1.43 mmol), and HOBt (63 mg, 0.57 mmol) were mixed in CH_2_Cl_2_ (1 mL) before EDC.HCl (118 mg, 0.62 mmol) was added. The reaction mixture was stirred at rt for 24 h before it was washed with HCl 2%, water, and brine. The organic layer was dried over MgSO_4_, filtered and reduced under vacuum. The crude residue was purified by flash chromatography over silica gel (CHCl_3_/MeOH 100:0 to 98:2) to afford **9a** (104 mg, 61%). *R*_f_ 0.8 (CHCl_3_/MeOH 8:2). ^1^H-NMR (CDCl_3_, 400 MHz) δ 6.47 (br., 1H), 5.69 (br., 1H), 4.94 (b., 1H), 3.81 (dq, *J* = 7.8, 6.4 Hz, 1H), 3.71 – 3.60 (m, 7H), 3.56 (dd, *J* = 5.6, 4.5 Hz, 2H), 3.44 (t, *J* = 4.9 Hz, 2H), 3.38 (dd, *J* = 5.5, 4.4 Hz, 2H), 2.17 (t, *J* = 7.4 Hz, 2H), 1.65 (p, *J* = 7.1 Hz, 2H), 1.52 – 1.23 (m, 6H), 1.11 (d, *J* = 6.5 Hz, 3H). ^13^C-NMR (CDCl_3_, 101 MHz) δ 173.10, 163.74, 70.44, 70.13, 70.07, 70.04, 56.08, 51.42, 50.65, 39.11, 35.95, 29.47, 28.63, 25.90, 25.21, 15.77. FT-IR (thin film) v 1650; 1697; 2107; 2863; 2933; 3270. HRMS (ESI) for C_16_H_31_N_6_O_4_^+^ [M+H]^+^, calcd 371.2401, found 371.2392 [24].

#### *N*-(17-Azido-3,6,9,12,15-pentaoxaheptadecyl)-6-((4*R*,5*S*)-5-methyl-2-oxoimidazolidin-4-yl)hexanamide (9b)

Compound **9b** (198 mg, 85%) was obtained from desthiobiotin and 17-azido-3,6,9,12,15-pentaoxaheptadecan-1-amine^4^ following the same protocol as for **9a**. *R*_f_ 0.65 (CHCl_3_/MeOH 8:2). ^1^H-NMR (CDCl_3_, 400 MHz) δ 6.55 (t, *J* = 5.6 Hz, 1H), 5.63 (s, 1H), 4.86 (s, 1H), 3.84 – 3.77 (m, 1H), 3.67 (t, *J* = 4.9 Hz, 2H), 3.67 –3.60 (m, 17H), 3.55 (dd, *J* = 5.6, 4.5 Hz, 2H), 3.42 (d, *J* = 5.4 Hz, 2H), 3.37 (d, *J* = 5.4 Hz, 2H), 2.18 (t, *J* = 7.4 Hz, 2H), 1.52 – 1.23 (m, 6H), 1.10 (d, *J* = 6.5 Hz, 3H). ^13^C-NMR (CDCl_3_, 101 MHz) δ 174.0, 163.6, 70.7 – 69.9 (br.), 56.01, 51.35, 50.64, 39.11, 35.92, 29.44, 28.61, 25.88, 25.16, 15.73. FT-IR (thin film) v 1697, 2100, 2863, 2930, 3292. HRMS (ESI) for C_22_H_43_N_6_O_7_^+^ [M+H]^+^, calcd 503.3188, found 503.3172 [25].

#### *N*-(2-(2-(2-Aminoethoxy)ethoxy)ethyl)-6-((4*R*,5*S*)-5-methyl-2-oxoimidazolidin-4-yl)hexanamide (10a). Azide 9a

(403 mg, 1.1 mmol) and Pd/C 10% (75 mg) in EtOH (10 mL) were vigorously stirred under a H_2_ atmosphere for 2 h. Catalyst was filtered off and volatile was removed under vacuum to yield compound **10a** (370 mg, 99%). *R*_f_ 0.2 (CH_2_Cl_2_/MeOH 8:2). ^1^H-NMR (CD_3_OD, 400 MHz) δ 3.91 – 3.75 (m, 1H), 3.75 – 3.66 (m,1H), 3.63 (m, 4H), 3.55 (t, *J* = 5.6 Hz, 3H), 3.36 (t, *J* = 5.6 Hz, 2H), 2.82 (t, *J* = 5.3 Hz, 2H), 2.21 (t, *J* = 7.5 Hz, 2H),1.65 – 1.57 (m, 4H), 1.53 – 1.42 (m, 2H), 1.40 – 1.29 (m, 2H), 1.11 (d, *J* = 6.4 Hz, 3H). ^13^C-NMR (CD_3_OD,101 MHz) δ 176.28, 164.62, 72.81, 71.33, 71.26, 70.62, 57.39, 52.70, 41.90, 40.27, 36.90, 30.74, 30.21, 27.15,26.81, 15.62. FT-IR (thin film) v 1691; 2864; 2933; 3269. HRMS (ESI) for C_16_H_33_N_4_O_4_^+^ [M+H]^+^, calcd 345.2496,found 345.2513.

#### *N*-(17-Amino-3,6,9,12,15-pentaoxaheptadecyl)-6-((4*R*,5*S*)-5-methyl-2-oxoimidazolidin-4-yl)hexanamide (10b)

Compound **10b** (210 mg, 99%) was obtained from **9b** following the same protocol as for **10a**. *R*_f_ 0.2 (CH_2_Cl_2_/MeOH 8:2). ^1^H-NMR (CD_3_OD, 400 MHz) δ 3.86 – 3.79 (m, 1H), 3.78 – 3.73 (m, 2H), 3.73 – 3.59 (m, 17H), 3.55 (t, *J* = 5.6 Hz, 2H), 3.36 (t, *J* = 5.6 Hz, 2H), 3.20 – 3.09 (m, 2H), 2.22 (t, *J* = 7.4 Hz, 3H), 1.64 (quin, *J* = 7.3 Hz, 2H), 1.56 – 1.21 (m, 6H), 1.11 (d, *J* = 6.4 Hz, 3H). ^13^C-NMR (CDCl_3_, 101 MHz) δ 176.3, 166.1, 79.59, 71.5 (b.), 71.42, 71.31, 71.26, 71.23, 71.14, 71.00, 70.65, 67.87, 57.36, 36.90, 30.73, 30.19, 27.13, 26.79, 15.64. FT-IR (thin film) v 1647, 1678,2869, 3269. HRMS (ESI) for C_22_H_45_N_4_O_7_^+^ [M+H]^+^, calcd 477.3283, found 477.3271.

#### 4-((3*R*,5*R*)-5-((5-Bromo-1-methyl-6-oxo-1,6-dihydropyridazin-4-yl)amino)-1-methylpiperidin-3-yl)-*N*-(2-(2-(2-(6-((4*R*,5*S*)-5-methyl-2-oxoimidazolidin-4-yl)hexanamido)ethoxy)ethoxy)ethyl)benzamide (1a)

A mixture of compound **8** (20 mg, 47.6 µmol), compound **10a** (24 mg, 69.6 µmol), EDC.HCl (37 mg, 193.6 µmol), HOBt (9 mg, 66.6 µmol), and Et_3_N (80 µL, 574 µmol) was stirred in anhydrous CH_2_Cl_2_ (4 mL) overnight at room temperature. The crude reaction mixture was washed with NaOH 1 N, brine, and the organic layer was dried over Na_2_SO_4_. Solvent was removed under vacuum and the residue was purified by flash chromatography over silica gel (CHCl_3_/MeOH 100:0 to 80:20) to afford **9a** (33 mg, 93%). *R*_f_ 0.6 (CHCl_3_/MeOH 8:2). ^1^H-NMR (CDCl_3_, 400 MHz) δ 7.77 (d, *J* = 8.3 Hz, 2H), 7.55 (s, 1H), 7.28 (d, *J* = 8.3 Hz, 2H), 6.98 (t, *J* = 5.5 Hz, 1H), 6.52 (t, *J* = 5.6 Hz, 1H), 5.78 (s, 1H), 4.86 (s, 1H), 4.65 (d, *J* = 8.7 Hz, 1H), 3.86 – 3.79 (m, 1H), 3.79 (q, *J* = 7.7 Hz, 1H), 3.74 (s, 3H), 3.69 – 3.59 (m, 7H), 3.54 (t, *J* = 5.3 Hz, 2H), 3.42 – 3.38 (m, 2H), 3.14 (d, *J* = 11.0 Hz, 1H), 3.04 (tt, *J* = 11.9, 3.7 Hz, 1H), 2.94 (d, *J* = 10.9 Hz, 1H), 2.36 (s, 3H), 2.28 (d, *J* = 12.4 Hz, 1H), 2.15 (t, *J* = 7.3 Hz, 2H), 2.05 (t, *J* = 11.2 Hz, 1H), 1.92 (t, *J* = 10.6 Hz, 1H), 1.62 (quin, *J* = 7.1 Hz, 2H), 1.54 – 1.22 (m, 7H), 1.09 (d, *J* = 6.4 Hz, 3H). ^13^C-NMR (CDCl_3_, 101 MHz) δ 173.10, 167.12, 163.68, 157.91, 145.76, 144.92, 133.11, 127.44, 127.21, 124.99, 100.05, 70.15, 70.12, 69.91, 69.81, 61.50, 61.04, 55.96, 51.38, 50.07, 45.84, 41.36, 40.34, 39.74, 39.08, 37.97, 35.90, 29.45, 28.53, 25.80, 25.16, 15.74. FT-IR (thin film) v 1604, 1693, 2930, 3292. HRMS (ESI) for C_34_H_52_BrN_8_O_6_^+^ [M+H]^+^, calcd 747.3188, found 747.3200.

#### 4-((3*R*,5*R*)-5-((5-Bromo-1-methyl-6-oxo-1,6-dihydropyridazin-4-yl)amino)-1-methylpiperidin-3-yl)-*N*-(24-((4*R*,5*S*)-5-methyl-2-oxoimidazolidin-4-yl)-19-oxo-3,6,9,12,15-pentaoxa-18-azatetracosyl)benzamide (1b)

Compound **1b** (43 mg, 56%) was obtained from **8** and **10b** following the same protocol as for **1a**. *R*_f_ 0.7 (CH_2_Cl_2_/MeOH 8:2). ^1^H-NMR (CDCl_3_, 400 MHz) δ 7.81 (d, *J* = 8.3 Hz, 2H), 7.56 (s, 1H), 7.28 (d, *J* = 12.5 Hz, 2H), 7.23 (t, *J* = 5.4 Hz, 1H), 6.67 (t, *J* = 5.6 Hz, 1H), 5.41 (s, 1H), 4.74 (s, 1H), 4.63 (d, *J* = 8.7 Hz, 1H), 3.92 –3.79 (m, 1H), 3.81 (quin, *J* = 6.7 Hz, 1H), 3.75 (s, 3H), 3.68 – 3.57 (m, 21H), 3.52 (t, *J* = 5.3 Hz, 2H), 3.48 – 3.35 (m, 2H), 3.17 (d, *J* = 11.0 Hz, 1H), 3.11 – 3.02 (m, 1H), 2.98 (d, *J* = 12.4 Hz, 1H), 2.38 (s, 3H), 2.28 (d, *J* = 12.1 Hz, 1H), 2.18 (t, *J* = 7.4 Hz, 1H), 2.07 (t, *J* = 11.2 Hz, 1H), 1.95 (t, *J* = 10.8 Hz, 1H), 1.63 (quin, *J* = 7.1 Hz, 2H), 1.58 – 1.15 (m, 7H), 1.10 (d, *J* = 6.5 Hz, 3H). ^13^C-NMR (CDCl_3_, 101 MHz) δ 173.11, 167.07, 163.45, 157.91, 145.41, 144.91, 133.30, 127.60, 127.10, 124.99, 100.12, 70.44, 70.42 (br.), 70.10, 70.03, 69.94, 69.92, 61.48, 60.94, 55.97, 51.34, 50.00, 45.77, 41.31, 40.36, 39.79, 39.11, 37.98, 35.99, 29.64, 29.48, 28.76, 25.94, 25.28, 15.74. FT-IR (thin film) v 1605, 1689, 2865, 2929, 3292. HRMS (ESI) for C_4_0H_64_BrN_8_O_9_^+^ [M+H]^+^, calcd 879.3974, found 879.3971.

### Cells preparation

K562 cells were cultured in Roswell Park Memorial Institute medium (RPMI) supplemented with 10% calf serum and 40 µg/ml gentamycin, at 37°C in 5% CO2 and 80 rpm rotative incubator. The cells have been grown into 6 L at 1.6 × 10^6^ C/ml.

HeLa cells were cultured in Minimal Essential Media Spinner Modification (S-MEM) supplemented with 7% newborn calf serum, 2 mM glutamine and 40 µg/ml gentamycin at 37°C in 5% CO_2_. The cells have been grown into 9 L at 1 × 10^6^ C/ml.

### Purification of SAGA/ATAC complexes

Three or four and half-liter cultures were centrifuged (1000 g for 10 min). All following steps were performed at 0-4°C. After centrifugation, the cell pellet was incubated 30 min on ice to equilibrate the transcription state, then was first washed with cold phosphate-buffered saline (PBS) followed by a wash with K75 buffer (10 mM HEPES pH 7.9; 75 mM KCl; 1.5 mM MgCl_2_; 1 mM DTT; 5 µg/ml pepstaine A and 3 µg/ml E64). Cell pellet was then resuspended in hypotonic buffer K0 (10 mM HEPES pH 7.9; 1.5 mM MgCl_2_; 1 mM DTT; 5 µg/ml pepstaine A; 3 µg/ml E64 and ROCHE protease inhibitor cocktail) and homogenized in a 100-ml B-Dounce homogenizer. Sucrose was added at a final concentration of 10% (w/v) and nuclei were pelleted for 28 min at 5,000g. Nuclear pellets were washed with sucrose buffer (10 mM HEPES pH 7.9; 10 mM KCl; 1.5 mM MgCl_2_; 10% sucrose w/v; 1 mM DTT; 5 µg/ml pepstaine A and 3 µg/ml E64). Nuclear pellets were resuspended in no-salt buffer (20 mM HEPES pH 7.9; 1.5 mM MgCl_2_; 25% glycerol w/v; 2 mM DTT; 5 µg/ml pepstaine A; 3 µg/ml E64 and ROCHE protease inhibitor cocktail) and homogenized in a 40-ml B-Dounce homogenizer. 0.3 M NaCl were added before a 30-min incubation with rotation. The nuclear extract was cleared by centrifugation (30,000g, 30 min) and flash-frozen in liquid nitrogen.

1.2% of polyethylene glycol 20,000 (PEG) and 5 mM MgCl_2_ were added to precipitate some remaining cellular organelles and membranes by centrifugation at 33,000g for 10 min. PEG 20,000 was then added at a 4.5% final concentration to precipitate large protein complexes with the same centrifugation step. The pellet was then resuspended in buffer A (20 mM HEPES pH8; 250 mM NaCl; 2mM MgCl_2_; 10% sucrose w/v; 0.5 mM DTT; 5 µg/ml pepstain A; 3 µg/ml E64; 0.05% Tween-20 and ROCHE protease inhibitor cocktail). The sample was mixed with Streptavidin Sepharose high performance beads (Cytiva) equilibrated in buffer A, and incubated with rotation for 4 hours to remove endogenous biotinylated proteins. In parallel, 35 µl of Streptavidin Sepharose high performance beads were incubated for two hours with 2.5 µl of the affinity ligand (10 mM stock solution), containing either the short (1a) or the long (1b) linker. The saturated beads were then washed twice with buffer A. The prepared sample and beads were mixed and incubated overnight. We used deliberately an amount of beads that would bind only 70% of the total mass of SAGA/ATAC in order to assure high purity of the sample. The beads were subjected to seven washes: three with buffer B (20 mM HEPES pH8; 250 mM NaCl; 2 mM MgCl_2_; 15% sucrose w/v; 0.5 mM DTT; 5 µg/ml pepstain A; 3 µg/ml E64; 0.05% Tween-20), one with buffer C (20 mM HEPES pH8; 250 mM NaCl; 2 mM MgCl_2_; 7.5% sucrose w/v; 5% trehalose; 2.5 µM β-mercaptoethanol (BME); 0.0025% DDM), and two with buffer D (20 mM HEPES pH8; 250 mM NaCl; 2 mM MgCl_2_; 5% trehalose; 2.5 µM BME; 0.002% dodecyl-maltoside (DDM)). The beads were finally eluted in buffer D without trehalose and containing 20 mM biotin. The eluate was used for both cryo-EM analysis and biochemical analysis.

We note that total signal derived from SAGA/ATAC subunits out of total PSM in the MS analysis is only 30% with the short linker, compared to 60% with the longer linker. Furthermore, the purification with the short linker-based conjugate produced overall 12 times less SAGA/ATAC than the purification with the longer version.

### Western blots for Figures 2 and Supplementary Figures 2 and 3

To allow comparison between the content of the nuclear extract to the sample following PEG precipitation we diluted the sample after PEG precipitation by the ratio between the volumes of the extract and the sample (roughly 20).

### Cryo-EM sample preparation and data acquisition

The sample was freshly cross-linked with 0.15% glutaraldehyde (GA). Three microliters of sample, concentrated at 0.15 mg/ml, were applied onto one side of a holey gold EM grid (UltrAuFoil R1.2/1.3 mesh) rendered hydrophilic by 90-s treatment in Fischione 1070 plasma cleaner operating at 34% power with a gas mixture of 80% argon and 20% oxygen. 1 µL of the sample was added on the other side of the grid. The grid was blotted on the 1-µL-side for 5 s and flash-frozen in liquid ethane using an EM GP2 Automatic Plunge Freezer at 6°C, 90% humidity.

### Data acquisition

The image dataset was acquired on a Titan Krios G4 microscope operating at 300 kV in nanoprobe mode using SerialEM for automated data collection. Movie frames were recorded on a Flacon 4i direct electron detector after a Selectris X energy filter using a 10 e-V slit width in zero-loss mode. Images were acquired at nominal magnification of 165,000 x yielding a pixel size of 0.729 Å.

### Image processing

The image dataset was pre-processed using cryoSPARC [26]. Particle coordinates were determined using crYOLO. Particle images were extracted using Relion 5 [27] with box size of 560 pixels rescaled to 256 pixels generating a pixel size of 1.595 Å. The dataset was analyzed in Relion 5 and cryoSPARC according to standard protocols. Briefly, three rounds of reference free 2D classification of the individual particle images were performed in cryoSPARC to remove images corresponding to contaminating or damaged particles and ice contaminations. References (3-D models) were generated by the ab initio 3-D reconstruction program of cryoSPARC. These structures were then used as references for 3D classification jobs in cryoSPARC and particles corresponding to high resolution 3-D classes were selected and used for non-uniform refinement. The selected particles were re-extracted in Relion 5 with box size of 560 pixels rescaled to 360 pixels generating a pixel size of 1.134 Å. These particles were refined in cryoSPARC and subjected to 3D classification in Relion 5 without alignment using various regularization parameter (T) values. Particles corresponding to high-resolution classes were used in the subsequent non-uniform refinement in cryoSPARC.

Focused refinements were carried out in cryoSPARC using masks covering the regions of interest created in ChimeraX [28].

### Model building

PDB structures 7KTS and 8H7G were used as starting models for building the atomic model. The model was refined in Phenix [29] by real-space refinement with secondary structure restrains and in Isolde. All display images were generated using ChimeraX.

### Reconstitution of histone octamers and preparation of nucleosome DNA

Octamers were reconstituted from individual *Xenopus leavis* (canonical) histones expressed as inclusion bodies according to the standard protocol [30, 31]. Widom-601 145 bp DNA was produced using a plasmid harboring 16 copies of this sequence as described by Dyer et al. [30].

### Nucleosome reconstitution

Nucleosomes with 145 bp Widom-601 positioning sequence were prepared according to NEB Dilution Assembly Protocol (E5350) (https://international.neb.com/protocols/2012/06/02/dilution-assembly-protocol-e5350) with some modifications as follows: 2.75 µM 145 bp Widom-601 DNA was mixed with 2.75 µM canonical histone octamers in a solution containing 2 M NaCl, 1 mM EDTA, 5 mM BME. The solution was incubated for 30 min at RT and then underwent serial dilutions down to 1.48 M, 1 M, 0.6 M, 0.25 M NaCl with buffer LowSalt (10 mM HepesKOH pH 8.0, 2.5 mM BME). After each dilution the solution was kept at RT for 30 min. In order to reduce the final NaCl concentration, nucleosomes were concentrated in 0.5 mL 100 kDa cutoff Amicon up to 100 µL, then diluted five times with buffer LowSalt. This step was repeated one more time. Finally, nucleosomes were concentrated to 3–4 µM and analyzed in a 5% native 0.2 x TBE polyacrylamide gel to ascertain the quality of the sample and absence of free DNA.

### Histone acetyltransferase assay

The reaction was performed in 20 mM HEPES pH 7.5, 50 mM NaCl, 2 mM MgCl_2_, 10% glycerol (v/v); 100 µg/ml BSA, 2.5 nM BME, 50 µM acetyl-CoA and 1 µM nucleosome. The reaction was started with the addition of 20 nM of SAGA/ATAC purified complexes. After 30 minutes of incubation at 37°C, the reaction was stopped by an addition of SDS loading buffer (Laemmli buffer) and heated at 95°C for 3 min. Proteins were then resolved by 15% SDS-PAGE and analyzed by western blot. The primary antibody used was against acetylated lysine (Cell Signaling Technology, 9441S).

### Deubiquitination assay

To test purified SAGA deubiquitinase activity, Ub-AMC from ENZO (ref. BML-SE211) was used. The reaction buffer is composed of 20 mM HEPES pH8, 150 mM Potassium Acetate, 5 mM MgCl_2_, 20% sucrose (w/v), 0.005% DDM, 0.1 mg/ml BSA and 230 nM Ub-AMC. The reaction was initiated with the addition of 20 nM of SAGA and was performed in 96-wells plate at room temperature in the dark. Measurements were done at 0, 5, 10, 15, 20, 25, 30, and 60 minutes in the PHERAstar^Plus^ plate reader with a wavelength excitation at 355 nm and emission at 460 nm. The emission intensity of Ub-AMC without enzyme was also measured as control.

## Acknowledgement

We acknowledge support from the Institut National de la Santé et de la Recherche Médicale (Inserm); the Centre National pour la Recherche Scientifique (CNRS); the Ligue Contre le Cancer; the University of Strasbourg Institute for Advanced Study (USIAS) for a fellowship to P.S. (IdEx Unistra); and the Agence Nationale de la Recherche grants to L.T. and G.P. (ANR-22-CE11-0013) and to the IGBMC (ANR-10-LABX-0030-INRT). This work of the Interdisciplinary Thematic Institute IMCBio+, as part of the ITI 2021-2028 program of the University of Strasbourg, CNRS and Inserm, was supported by IdEx Unistra (ANR-10-IDEX-0002), the SFRI-STRAT’US project (ANR-20-SFRI-0012) and EUR IMCBio (ANR-17-EURE-0023) under the framework of the France 2030 Program. We acknowledge the use of resources of the French Infrastructure for Integrated Structural Biology (FRISBI) (ANR-10-INBS-0005) and of Instruct-ERIC. The cryo-electron microscopes were co-financed by the European Regional Development Fund (ERDF), the Strasbourg Eurometropole, the Alsace Region, FRISBI and the ESR/EquipEx+ France-Cryo-EM (ANR-21-ESRE-0046).

## Data Availability

The experimental cryo-EM maps have been deposited in the Electron Microscopy Data Bank (EMBD) under accession codes EMD-53895, EMD-53896, EMD-53932, EMD-53934 and EMD-53935. A composite map was also deposited for SAGA (EMDB-53937). The model coordinates for SAGA derived from the composite maps have been deposited in the PDB database under the accession code 9RDK.

## References

1. Glinsky, G.V., O. Berezovska, and A.B. Glinskii, Microarray analysis identifies a death-from-cancer signature predicting therapy failure in patients with multiple types of cancer. J Clin Invest, 2005. 115(6): p. 1503–21.

2. El-Saafin, F., et al., SAGA-Dependent Histone H2Bub1 Deubiquitination Is Essential for Cellular Ubiquitin Balance during Embryonic Development. Int J Mol Sci, 2022. 23(13).

3. Bondy-Chorney, E., et al., Nonhistone targets of KAT2A and KAT2B implicated in cancer biology (1). Biochem Cell Biol, 2019. 97(1): p. 30–45.

4. Yayli, G., et al., ATAC and SAGA co-activator complexes utilize co-translational assembly, but their cellular localization properties and functions are distinct. Cell Rep, 2023. 42(9): p. 113099.

5. Herbst, D.A., et al., Structure of the human SAGA coactivator complex. Nat Struct Mol Biol, 2021. 28(12): p. 989–996.

6. Papai, G., et al., Structure of SAGA and mechanism of TBP deposition on gene promoters. Nature, 2020. 577(7792): p. 711–716.

7. Spedale, G., H.T. Timmers, and W.W. Pijnappel, ATAC-king the complexity of SAGA during evolution. Genes Dev, 2012. 26(6): p. 527–41.

8. Kalkat, M., et al., MYC Deregulation in Primary Human Cancers. Genes (Basel), 2017. 8(6).

9. Li, C., et al., Structure of the human TIP60-C histone exchange and acetyltransferase complex. Nature, 2024. 635(8039): p. 764–769.

10. Humphreys, P.G., et al., Discovery of a Potent, Cell Penetrant, and Selective p300/CBP-Associated Factor (PCAF)/General Control Nonderepressible 5 (GCN5) Bromodomain Chemical Probe. J Med Chem, 2017. 60(2): p. 695–709.

11. Bassi, Z.I., et al., Modulating PCAF/GCN5 Immune Cell Function through a PROTAC Approach. ACS Chem Biol, 2018. 13(10): p. 2862–2867.

12. Steijvoort B. F. V., et al., Remote Functionalization: Palladium-Catalyzed C5(sp3)-H Arylation of 1-Boc-3-aminopiperidine through the Use of a Bidentate Directing Group. ACS Catalysis, 2016: p. 4486–4490.

13. Morgan, M.T. and C. Wolberger, Competition Assay for Measuring Deubiquitinating Enzyme Substrate Affinity. Methods Mol Biol, 2018. 1844: p. 59–70.

14. Han, Y., et al., Architecture of the Saccharomyces cerevisiae SAGA transcription coactivator complex. EMBO J, 2014. 33(21): p. 2534–46.

15. Wu, P.Y. and F. Winston, Analysis of Spt7 function in the Saccharomyces cerevisiae SAGA coactivator complex. Mol Cell Biol, 2002. 22(15): p. 5367–79.

16. Cretu, C., et al., Molecular Architecture of SF3b and Structural Consequences of Its Cancer-Related Mutations. Mol Cell, 2016. 64(2): p. 307–319.

17. Haselbach, D., et al., Structure and Conformational Dynamics of the Human Spliceosomal B(act) Complex. Cell, 2018. 172(3): p. 454–464 e11.

18. Wang, H., et al., Structure of the transcription coactivator SAGA. Nature, 2020. 577(7792): p. 717–720.

19. Niewiadomska-Cimicka, A., A. Hache, and Y. Trottier, Gene Deregulation and Underlying Mechanisms in Spinocerebellar Ataxias With Polyglutamine Expansion. Front Neurosci, 2020. 14: p. 571.

20. Cortez, J.T., et al., CRISPR screen in regulatory T cells reveals modulators of Foxp3. Nature, 2020. 582(7812): p. 416–420.

21. Chen, J., N. Sawyer, and L. Regan, Protein-protein interactions: general trends in the relationship between binding affinity and interfacial buried surface area. Protein Sci, 2013. 22(4): p. 510–5.

22. Shao, W., et al., Prp5-Spt8/Spt3 interaction mediates a reciprocal coupling between splicing and transcription. Nucleic Acids Res, 2020. 48(11): p. 5799–5813.

23. Zhang, Z., et al., Structural insights into how Prp5 proofreads the pre-mRNA branch site. Nature, 2021. 596(7871): p. 296–300.

24. Klein, E., et al., New chemical tools for investigating human mitotic kinesin Eg5. Bioorg Med Chem, 2007. 15(19): p. 6474–88.

25. M. B. Andrus, T.M.T. E. P. Updegraff, Z. E. Sauna, S. V. Ambudkar, Synthesis and analysis of polyethylene glycol linked P-glycoprotein-specific homodimers based on (−)-stipiamide. Tetrahedron letters, 2001. 42(23): p. 3819–3822.

26. Punjani, A., et al., cryoSPARC: algorithms for rapid unsupervised cryo-EM structure determination. Nat Methods, 2017. 14(3): p. 290–296.

27. Kimanius, D., et al., New tools for automated cryo-EM single-particle analysis in RELION-4.0. Biochem J, 2021. 478(24): p. 4169–4185.

28. Goddard, T.D., et al., UCSF ChimeraX: Meeting modern challenges in visualization and analysis. Protein Sci, 2018. 27(1): p. 14–25.

29. Liebschner, D., et al., Macromolecular structure determination using X-rays, neutrons and electrons: recent developments in Phenix. Acta Crystallogr D Struct Biol, 2019. 75(Pt 10): p. 861–877.

30. Dyer, P.N., et al., Reconstitution of nucleosome core particles from recombinant histones and DNA. Methods Enzymol, 2004. 375: p. 23–44.

31. Luger, K., T.J. Rechsteiner, and T.J. Richmond, Expression and purification of recombinant histones and nucleosome reconstitution. Methods Mol Biol, 1999. 119: p. 1–16.

