## Supplementary Data for "Novel insights into the structure and evolution of the human SAGA complex by affinity-ligand purification"

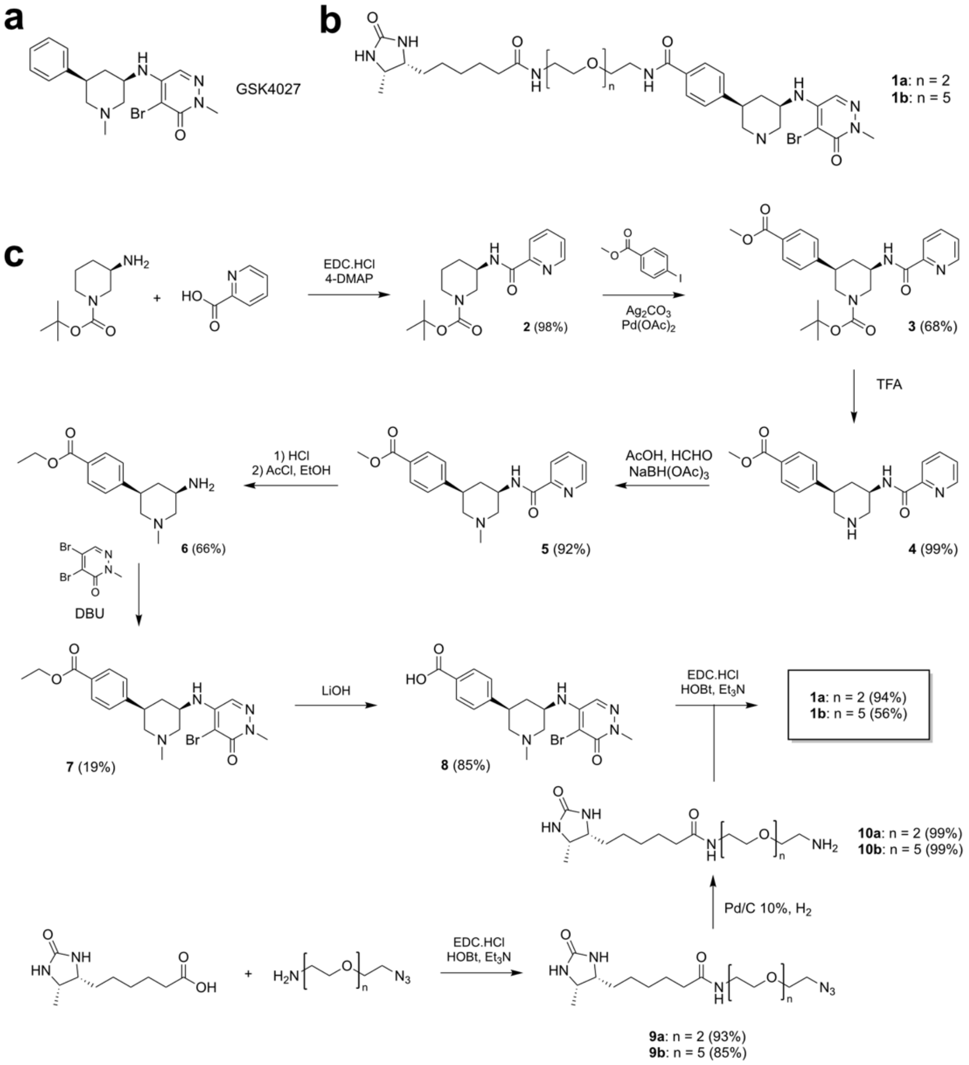


**Supplemental Figure 1: Synthesis of the affinity-ligands.**

**a,** structure of GSK4027. **b,** Structure of affinity ligands 1a and 1b. **c,** details of the synthetic route to prepare the compounds.

**a**

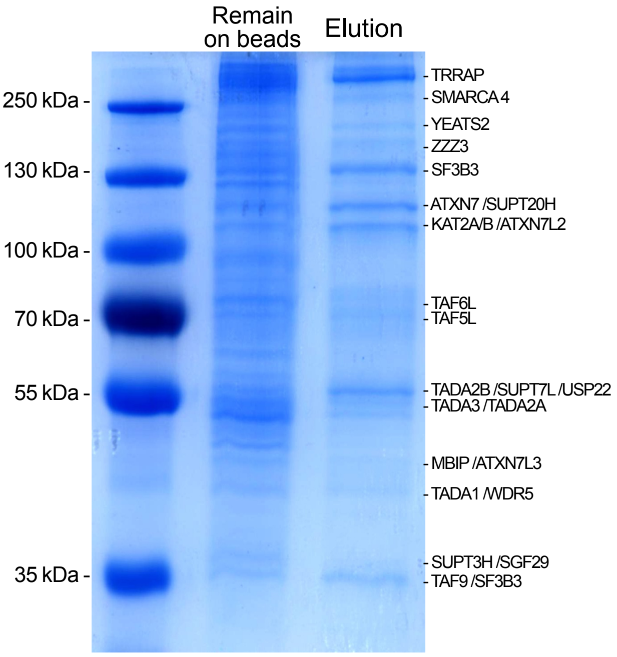


**b**

| Real position in MS list | Description | NSAF (PSM/AA) | Normalization (SUPT20H) |
|---|---|---|---|
| 1 | TRRAP | 0.91 | 2.2 |
| 2 | SF3B3 | 0.97 | 2.4 |
| 3 | BPTF | 0.23 | 0.6 |
| 4 | YEATS2 | 0.49 | 1.2 |
| 5 | KAT2A | 0.75 | 1.9 |
| 6 | TAF6L | 1.00 | 2.5 |
| 7 | TADA3 | 1.14 | 2.8 |
| 8 | SMARCA5 | 0.47 | 1.2 |
| 9 | TAF5L | 0.77 | 1.9 |
| 10 | BRD3 | 0.62 | 1.5 |
| 11 | ATXN7 | 0.42 | 1.0 |
| 12 | TADA2B | 0.88 | 2.2 |
| 13 | KAT2B | 0.44 | 1.1 |
| 14 | TAF9B | 1.42 | 3.5 |
| 15 | BRD4 | 0.25 | 0.6 |
| 16 | ZZZ3 | 0.36 | 0.9 |
| 17 | SUPT20H | 0.40 | 1.0 |
| 18 | TAF9 | 1.17 | 2.9 |
| 19 | DDX17 | 0.41 | 1.0 |
| 20 | ATXN7L2 | 0.40 | 1.0 |
| 22 | TADA1 | 0.79 | 1.9 |
| 28 | SGF29 | 0.72 | 1.8 |
| 31 | USP22 | 0.34 | 0.8 |
| 33 | MBIP | 0.51 | 1.3 |
| 36 | KAT14 | 0.21 | 0.5 |
| 37 | ATXN7L3 | 0.48 | 1.2 |
| 38 | SUPT7L | 0.39 | 1.0 |
| 44 | ATXN7L1 | 0.17 | 0.4 |
| 46 | TADA2A | 0.31 | 0.8 |
| 59 | SUPT3H | 0.34 | 0.8 |
| 77 | TAF10 | 0.39 | 1.0 |
| 79 | SF3B5 | 0.93 | 2.3 |
| 80 | WDR5 | 0.24 | 0.6 |
| 81 | TAF12 | 0.48 | 1.2 |
| 138 | ENY2 | 0.43 | 1.1 |
| 227 | DR1 | 0.13 | 0.3 |

**Supplemental Figure 2: Long linker purification on HeLa cells biochemical analysis**

**a,** Colloidal Coomassie blue stained SDS-PAGE of the purified SAGA/ATAC complexes. **b,** Proteomic analysis of the purified SAGA and ATAC complexes. For each identified protein the table shows the NSAF value calculated from the Peptide Spectrum Matches (PSM) divided by the number of amino acids, and the normalization of the NSAF value to SUPT20H as rough estimation of stoichiometry. SAGA subunits are colored purple, ATAC subunits are colored blue, purple bold represents the common subunits between SAGA and ATAC, and co-purified bromodomain containing proteins are in green.

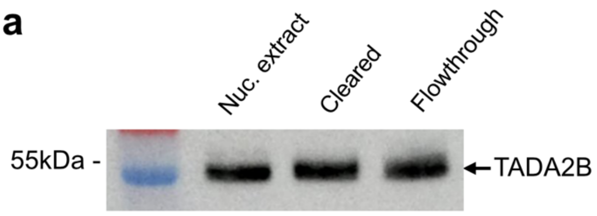


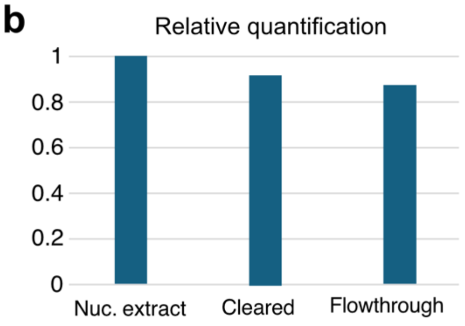


c

| Real position in MS list | Description | NSAF (PSM/AA) | NSAF norm. to SUTP20H |
|---|---|---|---|
| 1 | TRRAP | 1.09 | 1.5 |
| 2 | SF3B3 | 1.17 | 1.6 |
| **3** | **TADA3** | **1.89** | **2.5** |
| 4 | YEATS2 | 0.57 | 0.8 |
| **5** | **KAT2A** | **0.93** | **1.3** |
| **6** | **KAT2B** | **0.88** | **1.2** |
| 7 | TAF6L | 1.14 | 1.5 |
| 8 | TAF5L | 1.19 | 1.6 |
| 9 | BPTF | 0.22 | 0.3 |
| 10 | ATXN7L2 | 0.85 | 1.1 |
| 11 | SUPT20H | 0.74 | 1.0 |
| 12 | TADA2B | 1.32 | 1.8 |
| 13 | TAF9 | 2.01 | 2.7 |
| 14 | SMARCA5 | 0.44 | 0.6 |
| 15 | TAF9B | 1.82 | 2.4 |
| 16 | ZZZ3 | 0.47 | 0.6 |
| 17 | ATXN7 | 0.43 | 0.6 |
| 18 | DDX5 | 0.62 | 0.8 |
| 19 | TADA1 | 1.11 | 1.5 |
| 20 | DDX17 | 0.50 | 0.7 |
| **26** | **SGF29** | **1.04** | **1.4** |
| 32 | SUPT7L | 0.60 | 0.8 |
| 37 | KAT14 | 0.29 | 0.4 |
| 39 | SUPT3H | 0.69 | 0.9 |
| 40 | ATXN7L3 | 0.63 | 0.8 |
| 42 | USP22 | 0.41 | 0.6 |
| 44 | MBIP | 0.60 | 0.8 |
| 49 | TADA2A | 0.43 | 0.6 |
| 64 | ATXN7L1 | 0.18 | 0.2 |
| 79 | TAF12 | 0.81 | 1.1 |
| 102 | WDR5 | 0.29 | 0.4 |
| 109 | SF3B5 | 1.08 | 1.5 |
| 140 | TAF10 | 0.36 | 0.5 |
| 217 | ENY2 | 0.49 | 0.7 |
| 189 | DR1 | 0.20 | 0.3 |

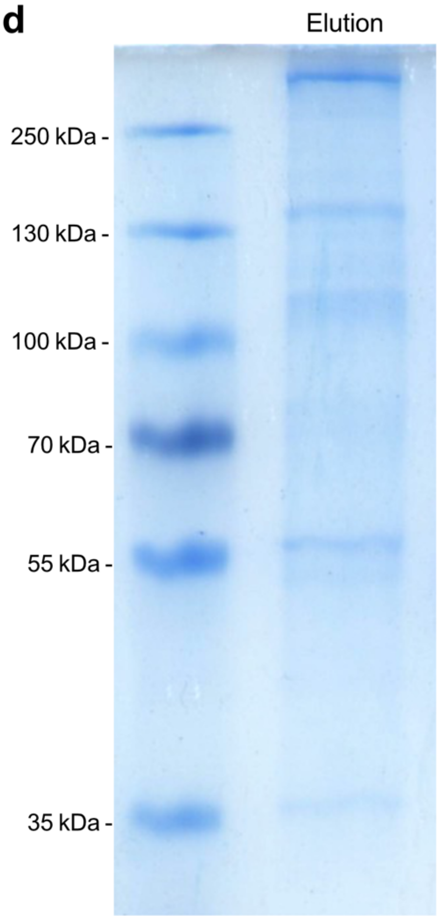


**Supplemental Figure 3: Purification of SAGA/ATAC using affinity-ligand with short linker**

**a,** Western blot analysis of different purification steps detecting the TADA2B subunit of SAGA. **b,** relative quantification of the detected bands in panel a. **c,** Proteomic analysis of the purified SAGA and ATAC complexes. For each identified protein the table shows the NSAF value calculated from the Peptide Spectrum Matches (PSM) divided by the number of amino acids, and the normalization of the NSAF value to SUPT20H as rough estimation of stoichiometry. SAGA subunits are colored purple, ATAC subunits are colored blue, purple bold represents the common subunits between SAGA and ATAC, and co-purified bromodomain containing proteins are in green. **d,** Colloidal Coomassie blue stained SDS-PAGE of the purified SAGA and ATAC complexes.

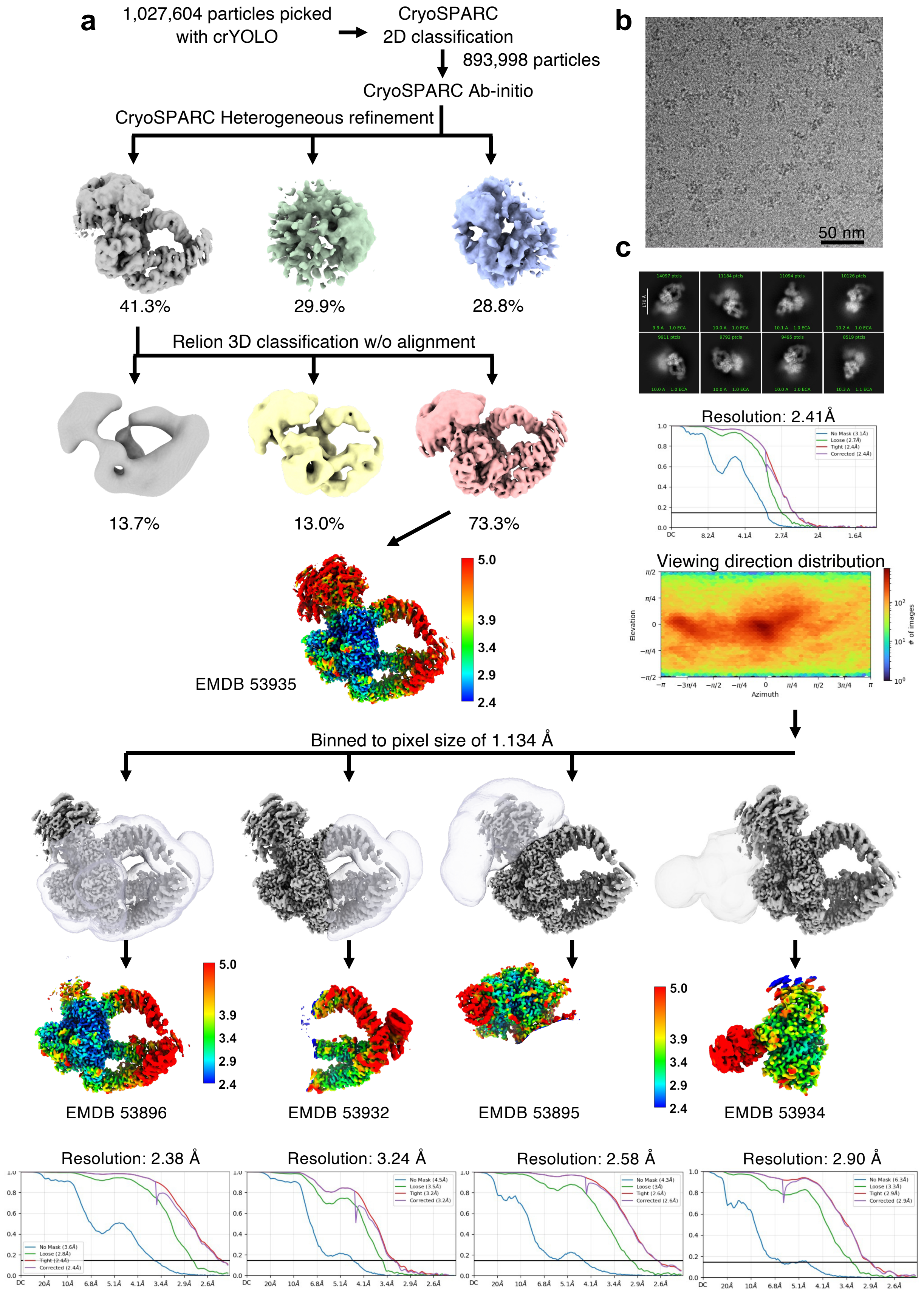


**Supplementary Figure 4: Cryo-EM data analysis strategy and resolution assessment**

**a,** Complete data processing scheme. Density map colored in rainbow represent the local resolution of the reconstructions. Hollow grey volumes represent the masks used for focused classification resolution in angstrom (Å) CryoSPARC v.4 were used to generate the 2.4 Å overall resolution map of human SAGA complex. RELION5 with Blush was used for classifications without alignment. **b,** Original micrograph. **c,** Two-dimensional class averages showing high-resolution structural features.

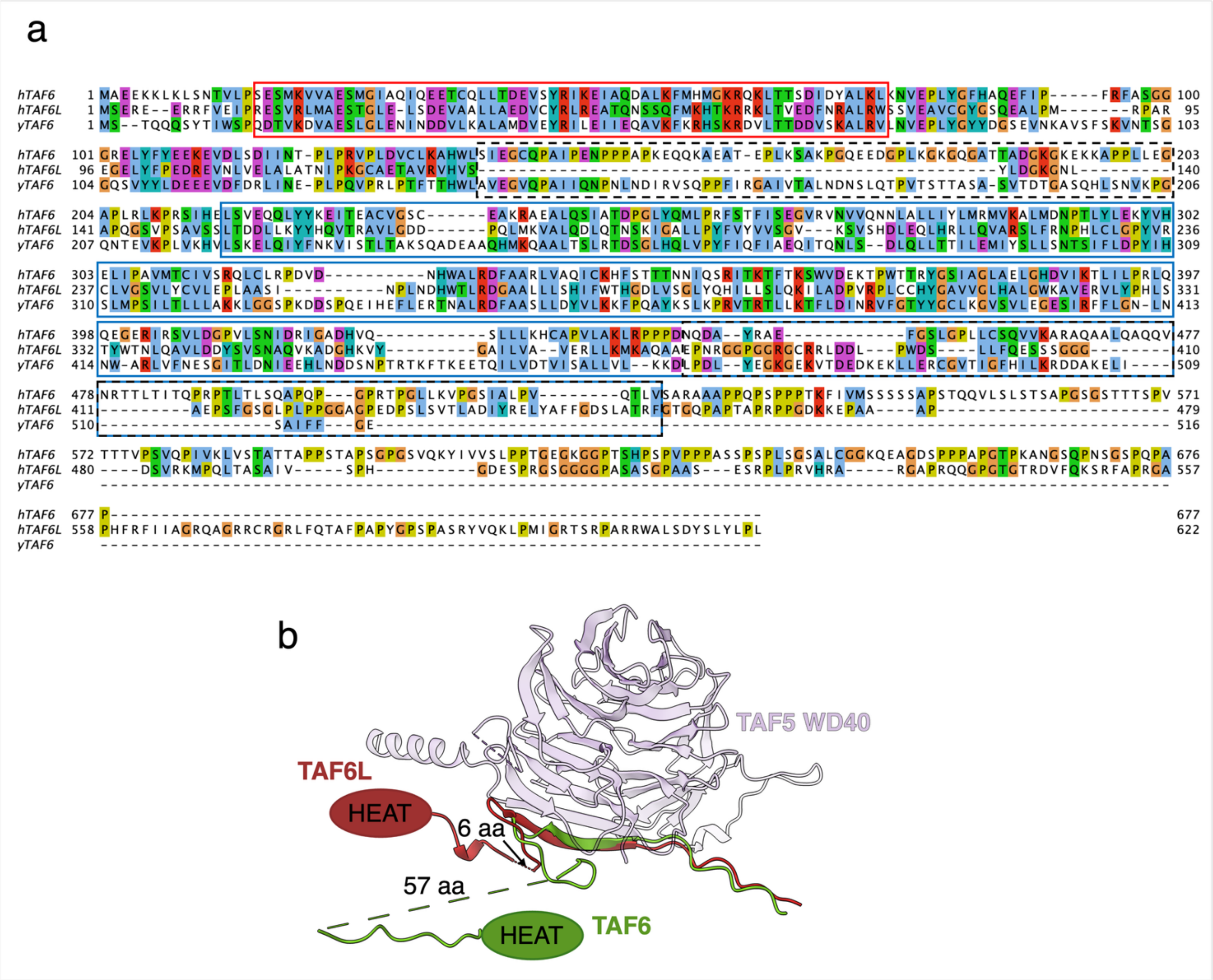


**Supplementary Figure 5: Sequence alignment of TAF6 and TAF6L**

**a,** Human TAF6, TAF6L and S. cerevisiae TAF6 were aligned using Clustal Omega. Boxes indicate structural features. Red: histone-fold domain; blue: HEAT repeat domain; dashed black: linker connecting histone fold with HEAT repeats; dashed black on blue: last two helices of HEAT repeats. **b,** TAF6L linker (in red), continuing the beta-strand complementing the the TAF5L WD40 repeat (light violet), is much shorter compared to the TAF6 linker (green). The TAF6L linker constrains the position of the HEAT repeats.

| | Overall (EMDB-53935) | TRRAP (EMDB-53896) | TRRAP-end (EMDB-53932) | Core (EMDB-53895) | SF3B (EMDB-53934) | Composite (EMDB-53937) (PDB 9RDK) |
|---|---|---|---|---|---|---|
| **Data collection and processing** | | | | | | |
| Magnification | 165,000 | 165,000 | 165,000 | 165,000 | 165,000 | 165,000 |
| Voltage (kV) | 300 | 300 | 300 | 300 | 300 | 300 |
| Electron exposure (e–/Å$^2$) | 40 | 40 | 40 | 40 | 40 | 40 |
| Defocus range (μm) | 1.2-3.5 | 1.2-3.5 | 1.2-3.5 | 1.2-3.5 | 1.2-3.5 | 1.2-3.5 |
| Pixel size (Å) | 0.729 | 0.729 | 0.729 | 0.729 | 0.729 | 0.729 |
| Symmetry imposed | C1 | C1 | C1 | C1 | C1 | C1 |
| Initial particle images (no.) | 1,027,610 | 1,027,610 | 1,027,610 | 1,027,610 | 1,027,610 | |
| Final particle images (no.) | 272,167 | 272,167 | 272,167 | 59,421 | 59,421 | |
| Map resolution (Å)<br>FSC threshold | 2.41<br>0.143 | 2.38<br>0.143 | 3.24<br>0.143 | 2.58<br>0.143 | 2.90<br>0.143 | |
| Map resolution range (Å) | 5-2.4 | 5-2.4 | 7-3.0 | 5-2.4 | 5-2.8 | 7-2.4 |
| **Refinement** | | | | | | |
| Initial model used (PDB code) | | | | | | 8H7G, 7KTR,7KTS |
| Model resolution (Å)<br>FSC threshold | | | | | | 2.3<br>0.143 |
| Map sharpening *B* factor (Å$^2$) | -58.8 | -47.6 | -100.4 | -40.5 | -62.4 | |
| **Model composition**<br>Non-hydrogen atoms<br>Protein residues<br>Ligands | | | | | | <br>53,996<br>6,734<br>0 |
| ***B* factors (Å$^2$)**<br>Protein | | | | | | <br>88.9 |
| **R.m.s. deviations**<br>Bond lengths (Å)<br>Bond angles (°) | | | | | | <br>0.005<br>1.064 |
| **Validation**<br>MolProbity score<br>Clashscore<br>Poor rotamers (%) | | | | | | <br>1.35<br>1.33<br>1.54 |
| **Ramachandran plot**<br>Favored (%)<br>Allowed (%)<br>Disallowed (%) | | | | | | <br>95.06<br>1.84<br>0.11 |

**Table S1. Cryo-EM data collection, refinement and validation statistics**
